# Genetic diversity and evolution of SARS-CoV-2 in Belgium during the first wave outbreak

**DOI:** 10.1101/2021.06.29.450330

**Authors:** Tony Wawina-Bokalanga, Joan Martí-Carreras, Bert Vanmechelen, Mandy Bloemen, Elke Wollants, Lies Laenen, Lize Cuypers, Kurt Beuselinck, Katrien Lagrou, Emmanuel André, Marc Van Ranst, Piet Maes

**Author notes:** Corresponding author; Herestraat 49 box 1040, BE3000 Leuven, Belgium. Contributed equally to this work (Tony Wawina-Bokalanga, Joan Martí-Carreras, Bert Vanmechelen).

## Abstract

SARS-CoV-2, the causative agent of COVID-19 was first detected in Belgium on 3rd February 2020, albeit the first epidemiological wave started in March and ended in June 2020. One year after the first epidemiological wave hit the country data analyses reveled the temporal and variant distribution of SARS-CoV-2 and its implication with Belgian epidemiological measures. In this study, 766 complete SARS-CoV-2 genomes of samples originating from the first epidemiological were sequenced to characterize the temporal and geographic distribution of the COVID-19 pandemic in Belgium through phylogenetic and variant analysis. Our analysis reveals the presence of the major circulating SARS-CoV-2 clades (G, GH and GR) and lineages circulating in Belgium at that time. Moreover, it contextualizes the density of SARS-CoV-2 cases over time with non-intervention measures taken to prevent the spread of SARS-CoV-2 in Belgium, specific international case imports and the functional implications of the most representative non-synonymous mutations present in Belgium between February to June 2020.

## Introduction

The first cases of coronavirus disease 2019 (COVID-19) were detected in December 2019 in Wuhan (China). Shortly thereafter, the causative agent was identified as SARS-CoV-2^1^. SARS-CoV-2 has spread worldwide in a short period of time, unlike the *Betacoronavirus* members SARS-CoV-1 (2002-2003) and MERS-CoV (2012 and 2015), whose outbreaks could be better contained, limiting international spread and averting global transmission^2–4^. Based on publicly available data, the case fatality rate of SARS-CoV-2 is estimated to be around 3% (https://www.worldometers.info/coronavirus/), lower than SARS-CoV-1 and MERS-CoV, with 9.6% and 34.3%, respectively ^5^, although the true infection fatality rate is likely lower, as many asymptomatic or mildly symptomatic cases remain undiagnosed^6^. Nonetheless, there is an urgent need for effective treatments, as the global death toll has already surpassed 3.7 million confirmed deaths (WHO coronavirus dashboard, 12/06/2021), and measures taken to control the spread of the virus have significantly impacted social and economic activity world-wide.

SARS-CoV-2 is a non-segmented, positive single-strand RNA (+ssRNA) virus with a genome length of around 30,000 nucleotides. The genome is organized into 12 genes that code for structural and non-structural proteins^5^. Structural proteins are the Spike (S), Envelope (E), Membrane (M) and Nucleocapsid (N) proteins. Non-structural proteins are all generated from the ORF1ab polyprotein, including the core proteins Nsp12 (RNA synthesis), Nsp7 and Nsp8^7^. Additionally, at least 5 accessory proteins are encoded alongside the structural proteins: ORF3a (putative apoptotic factor), ORF6 (putative IFN-1 antagonist), ORF7a (putative leukocyte modulator) ORF8 (putative immunomodulator), and ORF10 (unknown function). Recent estimates situate the mutation rate of SARS-CoV-2 between 0.8 × 10^−3^ and 1.12 × 10^−3^ substitutions/site/year^8,9^, equating to 2 to 2.8 substitutions/month. As per the writing of this manuscript, SARS-CoV-2 mutations have been organized dynamically in three large clades, based on non-synonymous substitutions^10^: (i) clade G (D614G in the Spike), (ii) clade V (G251 in ORF3a) and (iii) clade S (L84S in ORF8)^11^. The Spike mutation at D614G has been recently linked to an increased virus production in host cells and seems to be the predominant mutation in European clades since March 2020^12^. Genetic diversity analyses currently play an important role in improving our understanding of SARS-CoV-2. Complete genome sequences shared through the Global Initiative on Sharing All Influenza Data (GISAID) have been, and still are, valuable to monitor and contain the pandemic. Additionally, this unprecedented international cooperation has allowed to rapidly evaluate the viral origin and genomic diversity of SARS-CoV-2^13^ based on sequence similarity. Likewise, the availability of sufficient and diverse genome sequences to capture the variability of SARS-CoV-2 may allow estimations of which sets of antiviral drugs are most likely to be repurposed for this virus^13^. Changes in infection and mortality rates are influenced by SARS-CoV-2 genetic variation^14^ and host genetics^15^.

In Belgium, the first confirmed case of SARS-CoV-2 infection was reported on 3th of February 2020 (2020 week 5), from an asymptomatic individual who was part of a quarantined group of 10 travelers, epatriated from Wuhan to Brussels^16^. During week 8 of 2020, the Belgian government started engaging in travel regulations from and into China, trying to minimize the possible SARS-CoV-2 introduction events into the country. Still, borders were not closed until week 11, thus several entry events occurred prior to strict border regulations. Later, the Belgian government attributed part of these introduction events to returning travelers from the Carnival break (Figure 1). In March 2020, the rapidly growing number of confirmed cases alarmed the government, who decided to take restrictive measures to reduce the spread of SARS-CoV-2 into and inside the country. By week 10, the federal government limited indoor activity nationwide by closing bars and restaurants and prohibiting sportive and cultural activities (lockdown). From week 11 to week 18, strict social distancing measures were applied, and all non-essential travel and gatherings were suspended. Strict isolation policy during this period, which included the Easter break (weeks 15-16, Figure 1) were key to a substantial reduction of cases. Part of the lockdown measures were alleviated at week 18, although most public establishments were only allowed to re-open to the public in week 23 (Figure 1). Positive cases steadily increased until mid April (2020 week 15), after which case numbers decreased until mid June, following several intervention measures (Figure 1, Suppl. Table 1). During this period, 59.242 COVID-19 cases were reported. For 776 of these cases, the complete SARS-CoV-2 genome has been sequenced (1.3% of positive cases, Suppl. Table 1), including the first re-infection case in Belgium and one of the first cases world-wide^17^. In this study, we performed phylogenetic and mutational analysis of SARS-CoV-2 genome sequences obtained during the first wave outbreak across different provinces in Belgium before the establishment of a Belgian consortia for genomic surveillance of SARS-CoV-2.

**Figure 1.**
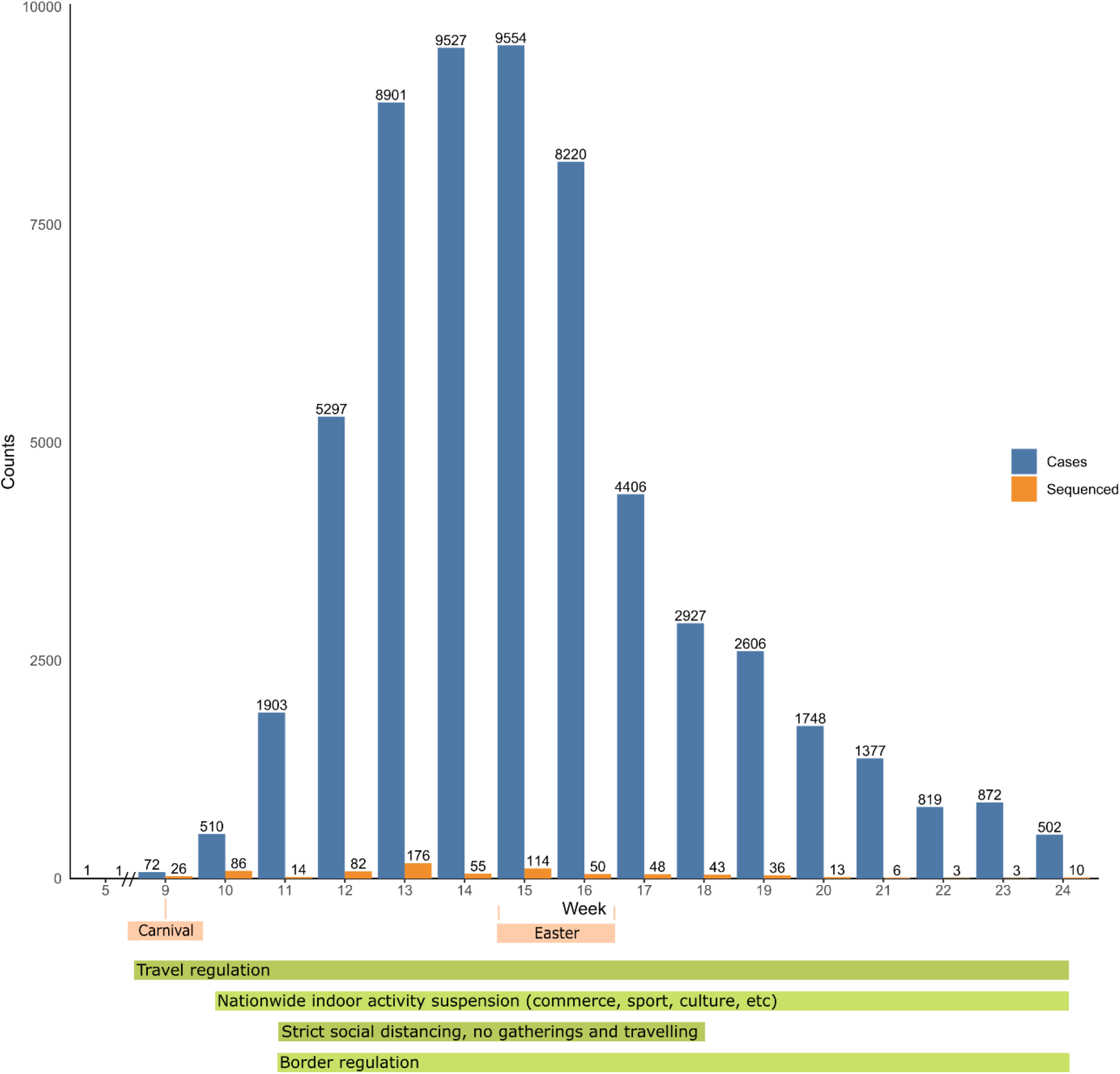
Progression of SARS-CoV-2 reported positive cases (blue) in relation to the sequencing effort (orange) during the first wave in Belgium (https://epistat.wiv-isp.be/covid/). At the bottom, a timeline is presented with school holidays (a week or more, in salmon) and key preventive measures taken by the Belgian federal government to stop the spread of SARS-CoV-2 (green).

## Material and Methods

### Sample collection

Samples were obtained through University Hospitals Leuven, the Belgian National Reference Center (NRC) for Respiratory Pathogens. Different types of samples including oro- and nasopharyngeal swabs, broncho-alveolar lavages and endotracheal aspirates were collected and tested for SARS-CoV-2 by Reverse Transcription qPCR at the NRC. Samples were randomly selected from a pseudo-anonymized database considering viral load (Ct < 20) and spread among collection dates and municipalities. Samples with low titulation coming from low-positivity regions or time periods were also considered in the study.

### RNA extraction and sequencing

RNA extraction was performed with 140 μl of swab transporting medium using the QIAampViral RNA extraction kit (ref. 52904, Qiagen). SARS-CoV-2 genomes were amplified following the ARTIC network protocol V1 and V2^16^. Briefly, reverse-transcription with SuperScript IV (ref. 18090010, Invitrogen) was performed on SARS-CoV-2 positive RNA extracts. Multiplex PCR with specific primer pools, tiling the complete SARS-CoV-2 genome, was performed on cDNA using Q5 Hot Start High-Fidelity DNA polymerase (ref. M0493S, New England Biolabs). Amplicons were immediately cleaned up with AMPure XP beads (New England Biolabs, ref. A63880) and libraries were prepared using the ligation sequencing kit (SQK-LSK109) from Oxford Nanopore Technologies with the modifications detailed in the ARTIC protocol. Libraries were quantified using QUBIT 1X dsDNA HS Assay Kit (ref. Q33231, Invitrogen). Sequencing was performed on MinION and GridION platforms using MinKNOW’s (v19.12) built-in basecalling, demultiplexing and adapter trimming (dual-barcode detection at 60% barcode sequence identity).

### Data collection and phylogenetic analysis

Sequencing runs were processed using the ARTIC analysis pipeline and custom scripts. Sequence metadata was collected and complemented to their respective GISAID records. SARS-CoV-2 lineages were derived using the Pangolin tool (https://cov-lineages.org/pangolin.html, github.com/cov-lineages/pangolin)^10^. Epidemiological data was obtained from Sciensano (the Belgian federal institute for health, https://epistat.wiv-isp.be/covid/). Sequences were aligned with MAFFT^18^, SNPs were collected with snp-sites^19^ and annotated with VCF-annotator (https://github.com/rpetit3/vcf-annotator). Data analysis was performed in R v3.5^20^ and Tidyverse^21^. Phylogenetic analysis were conducted with IQTREE^22^, TimeTree^23^ and JModelTest^24^ using the non-redundant set of Belgian SARS-CoV-2 genomes (528 sequences) to generate a time-scaled Maximum Likelihood phylogenetic tree (ML). Full-genome sequences are publicly available in GISAID with accession IDs in Suppl. File 1 and their Pangolin clade assignation in Suppl. Figure 1 and Suppl. Table 2.

## Results

### Geographic distribution and sequencing coverage

The first wave of the SARS-CoV-2 pandemic hit Belgium in March 2020, although the first imported case was already detected in the first days of February 2020. Throughout the first wave, samples were transferred from across Belgium (Figure 2) through the NRC for Respiratory Pathogens. Sequencing covered all 10 Belgian provinces and the Brussels-Capital Region, with the exception of the sparsely populated Luxembourg province (representing <2.5% of the Belgian population). Overall sequence density (SARS-CoV-2 genome sequence counts per town and province) is wide-spread, with the provinces of Limburg, Vlaams-Brabant and Brussels being slightly overrepresented due to the geographic proximity of these provinces to the location of the NRC. Sequencing rate (proportion of sequenced cases per total cases reported) was ~1% throughout most of the period, with the exception of the first weeks of the pandemic, in which the number of detected cases was low and hence a higher sequencing rate of 36% - 100% (Suppl. Table 1).

**Figure 2.**
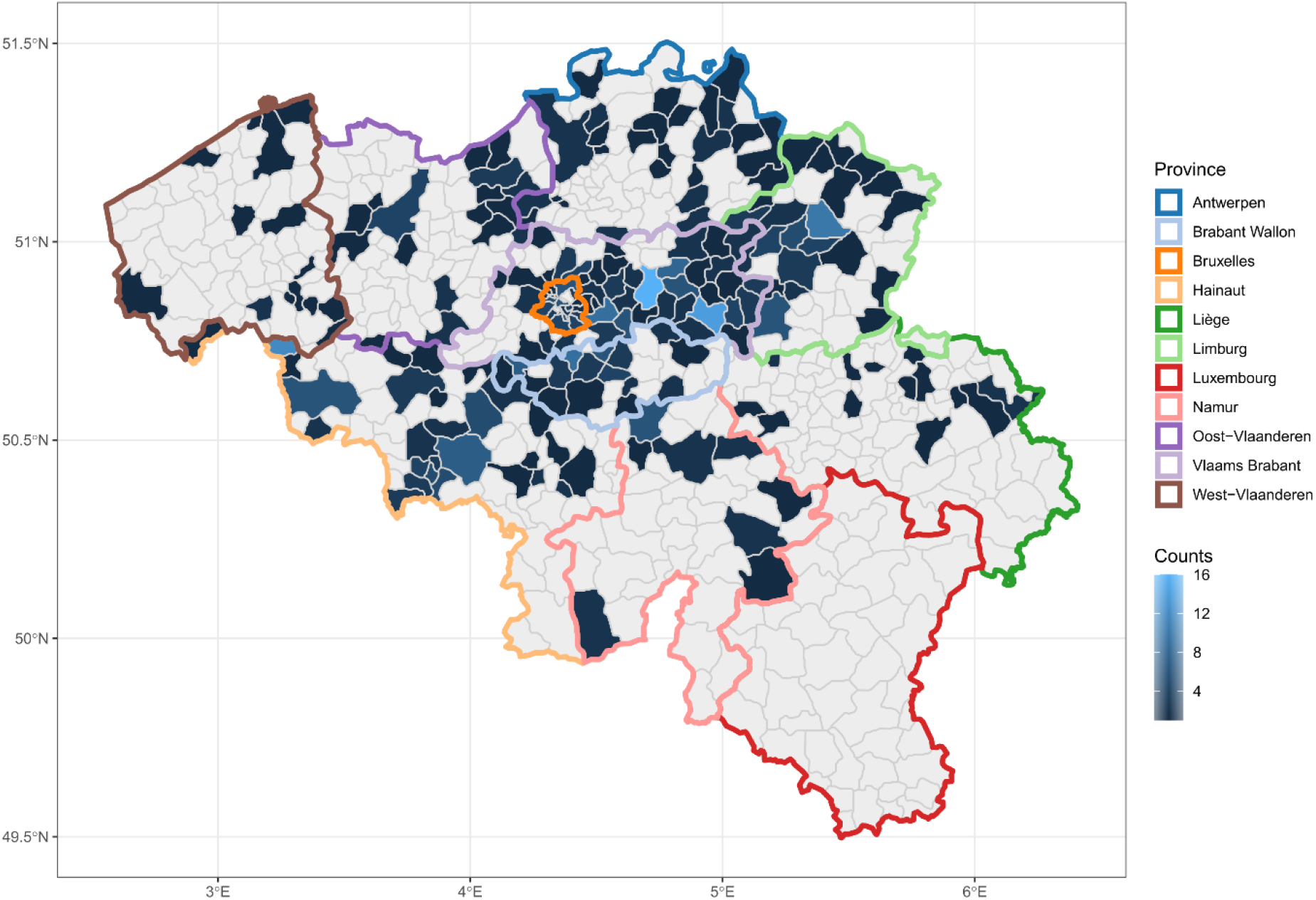
Geographical spread of SARS-CoV-2 genomes sequenced in Belgium during the first wave (March to 15 June 2020). Axes represent the cartesian coordinates of Belgium and its municipalities. Belgium is divided by municipalities (grey rim), colored in blue gradients depending on the number of SARS-CoV-2 genomes sequenced from that municipality (light grey for 0). Belgian provinces (10) and the Brussels-Capital Region (1) delimitations are highlighted by colour.

### Temporal distribution and phylogeny

The temporal distribution of complete SARS-CoV-2 genomes can be observed with a time-scaled maximum likelihood-tree (ML), in which sampling dates (tip dates) re-calibrate the phylogenetic distance between sequences (Figure 3), as previously used elsewhere ^16^. Most of the sequenced cases correspond to ‘domestic’ spread of the virus (within the country), as most of the tips in the phylogenetic tree tend to cluster at short distance with other tips. Tips with long branch lengths that occupy single clusters without neighbouring tips may represent international re-introductions or undersampled ‘domestic’ events. The latter becomes more relevant as the number of available sequences decreases over time (especially during May and June of 2020). Interestingly, 4 tips are clustered apart from the rest of the phylogenetic tree, indicating strong divergence from the rest of the tree (Figure 3, dashed rectangle). These cases correspond to direct internationally imported events (i) before non-essential international travel was banned (original case in February and a secondary case in March), or (ii) despite border regulations, as the sequences represented by the tips in April and May do not cluster with the rest of the tree, indicating a direct import event (after border regulations were enforced).

**Figure 3.**
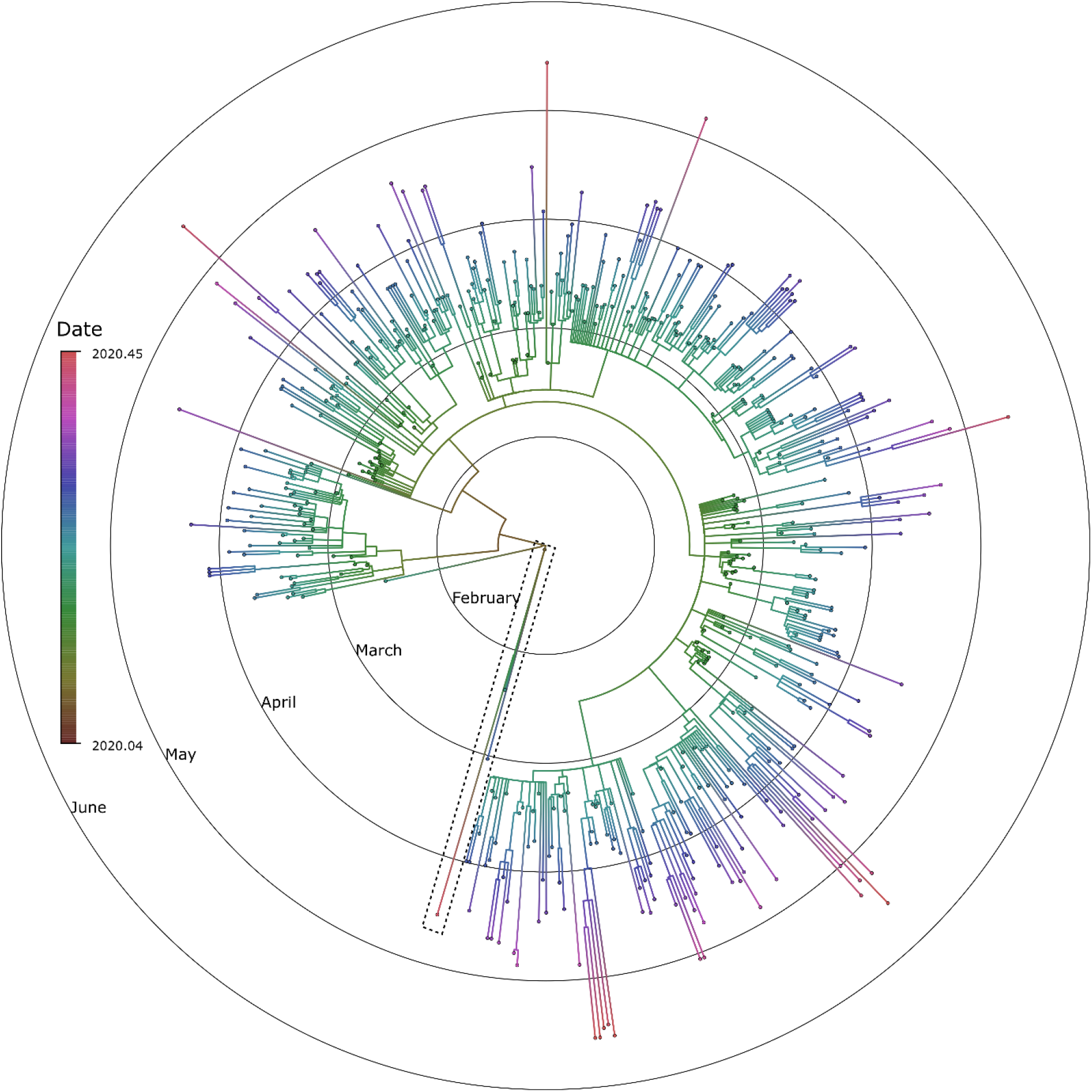
Phylogenetic distribution of SARS-CoV-2 genomes sequenced in Belgium during the first wave (February to 15 June 2020). Concentric circles represent months (February to June 2020). A tip-dated Maximum Likelihood phylogenetic tree (ML) represents the phylogenetic distance between the SARS-CoV-2 genomes re-scaled using sampling dates (using TimeTree^23^). The ML tree was built using the GTR + I phylogenetic model (inferred using JModelTest^24^). Branches and tips are coloured based on a temporal gradient (legend at the right side), where red/brown corresponds to the most ancestral state and purple/pink to the most recent. Dashed rectangle marks 4 sequences that cluster apart from the tree at different times, indicating imported events. Node ages have been removed from the tree for visualization purposes.

Tip density in the ML tree is not constant over time, in part due to sequence downsampling (merge sequences at 100% identity and same collection date), sequencing availability (access to sample for sequencing) and differences in COVID-19 positivity rates (Figure 1). The highest density is found throughout April, following the trend of COVID-19 positive cases and SARS-CoV-2 sequenced genomes shown in Figure 1. As hinted previously, intervention measures applied in mid March (week 11, Figure 1) reduced the transmission of SARS-CoV-2 in the Belgian population, reducing reported positive cases from May onwards. Correspondingly, a declining tip density can be observed for May-June in the ML tree. Despite higher sequencing rates would have been desirable for the study; sample accessibility and reagent discontinuity forbade a deeper genomic surveillance on the Belgian population. Nevertheless, as shown in Dellicour *et al*., the number of cases sequenced and its dispersion between March – June provide sufficient resolution to identify and trace SARS-CoV-2 lineages through time ^16^.

### Isolate classification and distribution of relevant mutations

SARS-CoV-2 acquires 2-3 nucleotide mutations per month (evolutionary rate 0.8 × 10^−3^ and 1.12 × 10^−3^ substitutions/site/year^8,9^). The acquisition of genetic changes allows the different genome sequences to be classified into groups. Currently, there are 2 widely used SARS-CoV-2 classifications based on: (i) specific amino acid changes (as found in GISAID, and called *Clade* hereafter or CoVariants in Nextclade ^25^) and (ii) a dynamic phylogenetic classification ^10^(conducted through the Pangolin classification tool, referred to as *Lineage* hereafter). SARS-CoV-2 clades are further subdivided into 7 different groups, depending on their mutations: L (C241, C3037, A23403, C8782, G11083, G26144, T28144), G (C241T, C3037T, A23403G nucleotide mutations and the aminoacid change D614G in the spike protein), GH (G clade mutations plus G25563T and the aminoacid change Q57H in NS3), GR (G clade mutations plus G28882A and the aminoacid change G204R in the nucleoprotein), S (C8782T, T28144C and the aminoacid change L84S in NS8), V (G11083T, G26144T and the amino acid changes L37F in NSP6 and G251V in NS3) and O (others). The lineage classification of SARS-CoV-2 makes use of Maximum-likelihood phylogenies of globally available viral sequences to dynamically name the branches of the tree with an alphanumeric system (defining first a major lineage, A or B, and then numbering the successive branching events). As shown by Alm *et al*., there is a correspondence between SARS-CoV-2 classification between GISAID clades, Pangolin lineages and the NextStrain classification scheme (not elaborated upon)^26^.

Distribution of clades and lineages of SARS-CoV-2 during the first epidemic wave of SARS-CoV-2 in Belgium strongly resembles the European distribution reported by Alm *et al*., with the G (Figure 4, blue) and GR (red) clades being the most prominent. GH (orange) emerges with an intermediate frequency, slowly diminishing towards the end. Together, the G clades, being the most prevalent ones, are likely responsible for most of the ‘domestic’ transmission cases. Clades L, S and V are seldom found during these initial 5 months of the SARS-CoV-2 epidemic in Belgium and were likely linked to direct import cases and their direct spread sphere. Specifically, clade S (the most ancestral SARS-CoV-2 clade) is the least prevalent, being detected only 4 times, each time linked to international travel and as seen in Figure 2, forming a clearly distinct clade directly rooted to the initial spreading events. Regarding lineages, the largest contributors were B.1 (G clade), B.1.1 (GR clade) and B.1.5 (G clade), especially in March and April (Table 1 and Suppl. Figure 1, respectively). Overall, more than 42 different lineages were detected over the course of 5 months, 40 representing B-lineages and 2 representing A-lineages (A and A.2).

**Figure 4.**
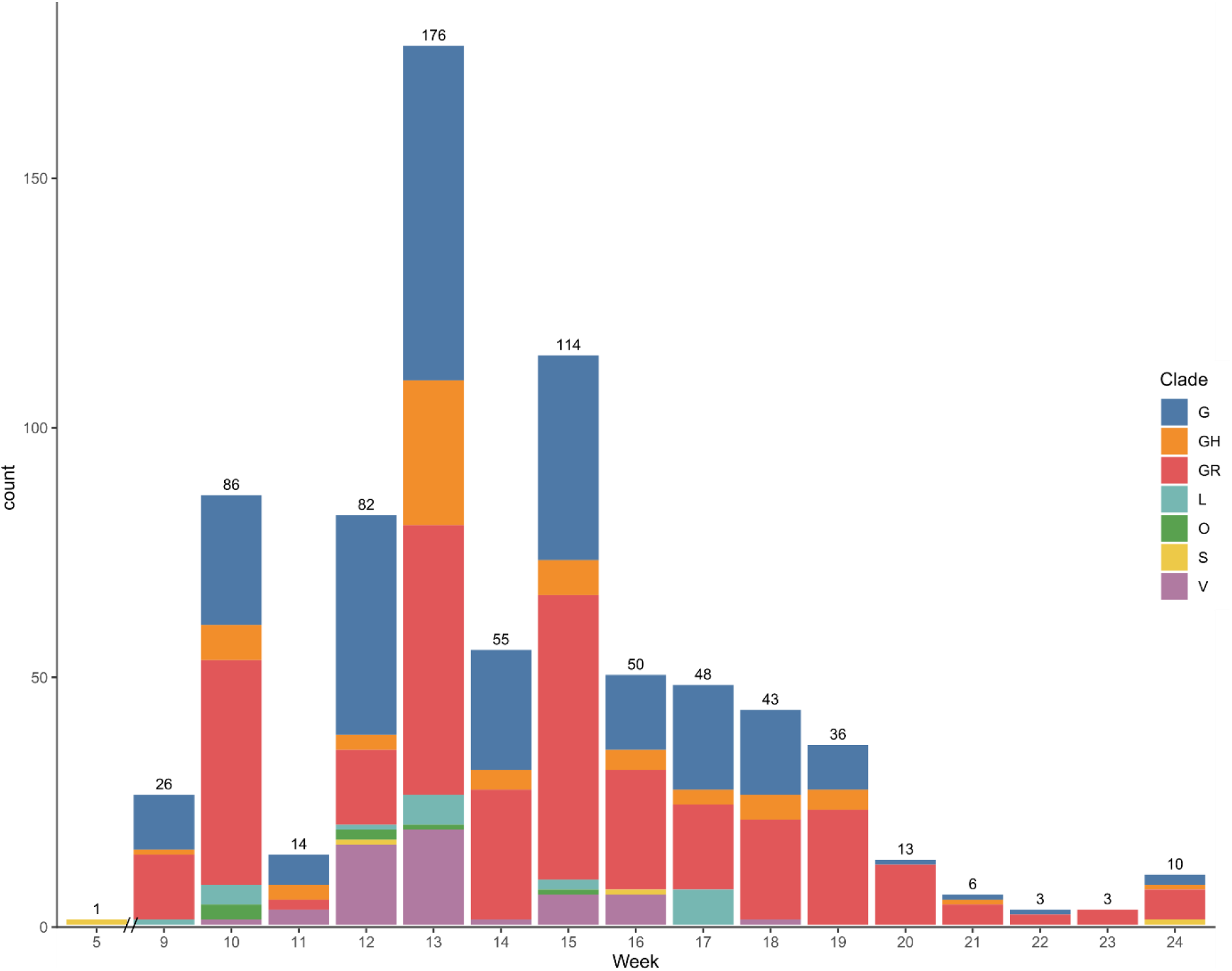
Clade classification of SARS-CoV-2 genomes during the first epidemic wave in Belgium. Number of sequences (y-axis as counts) is distributed during the weeks of February to June (x-axis). Sequences are classified into clades, following the GISAID classification (G, GH, GR, L, O, S, V), as shown in the legend. Total number of SARS-CoV-2 genomes per week are marked on top of each individual bar.

**Table 1.**
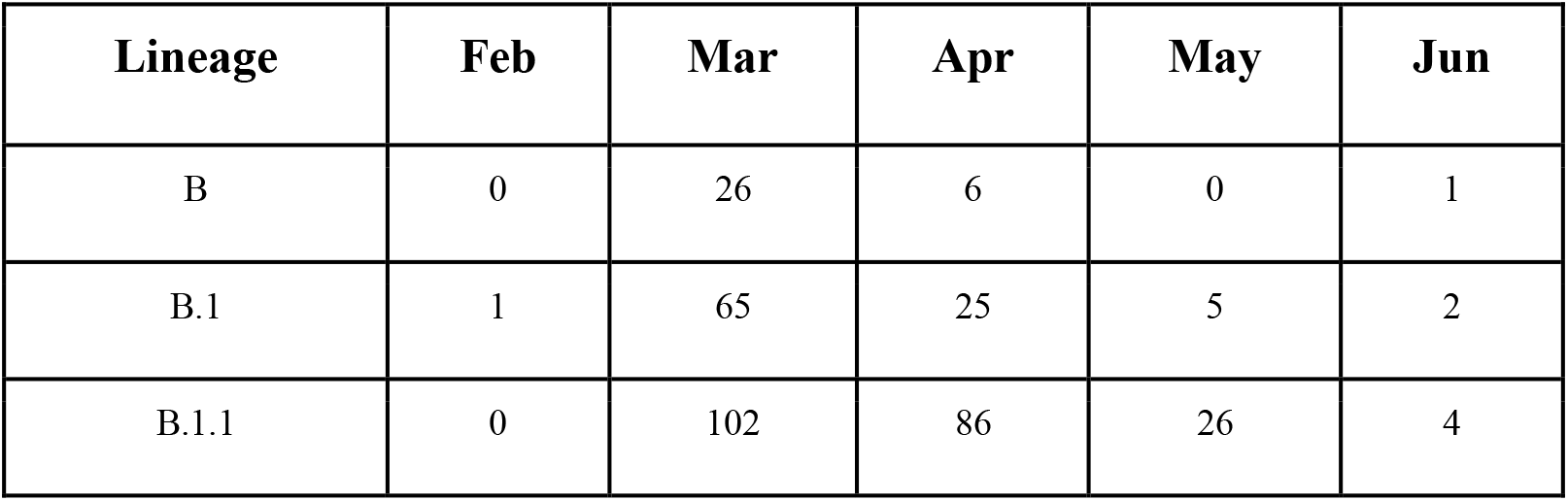
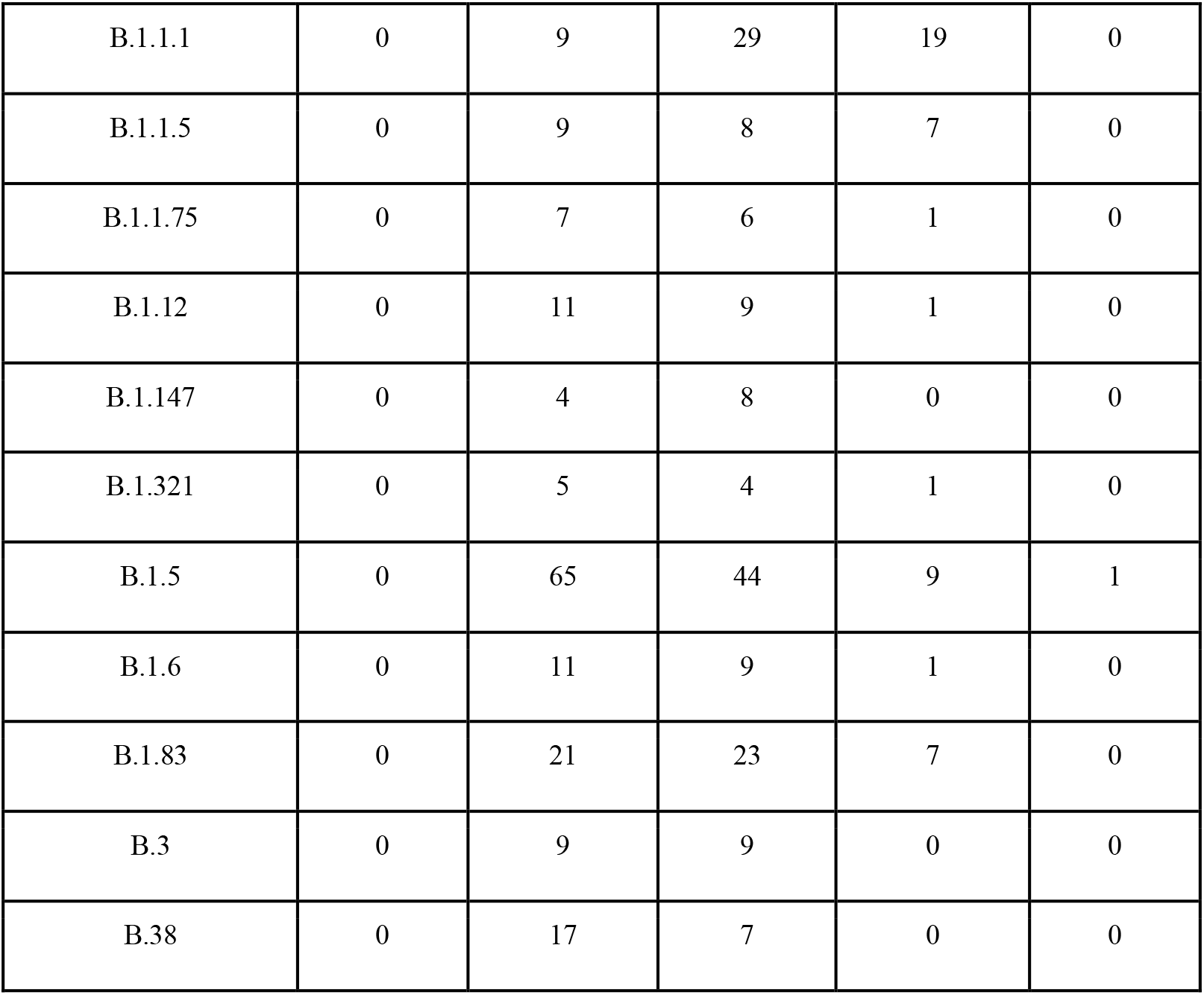
Most frequent (>10) SARS-CoV-2 lineages detected during the first COVID-19 wave in Belgium by month using the Pangolin tool. A complete summary is available in Suppl. Table 2 and Suppl. Figure 1.

Non-synonymous mutations were distributed across the entire SARS-CoV-2 genome (Figure 5). Certain amino acid changes were found to be more frequent in the complete dataset. An amino acid change was deemed frequent when it was present in at least 10% of sequences of the dataset (n = 776, 77 sequence count threshold). We observed five frequent mutations: L3606F (ORF1a, NSP6), T5020I (ORF1ab, NSP12), I65256T (ORF1ab, NSP15), D614G (S gene), R203K and G204K (N gene). The prevalence of some of these mutations can be traced back directly to specific clades, as some clades are defined by specific amino acid changes. Examples are D614G, which is linked to clade G and its derivatives (GH and GR) or G204K, which is present in clade GR. Other mutations that are characteristic of specific clades, such as Q57H in the NS3 (GH clade), are present in the dataset but at frequencies below 10%.

**Figure 5.**
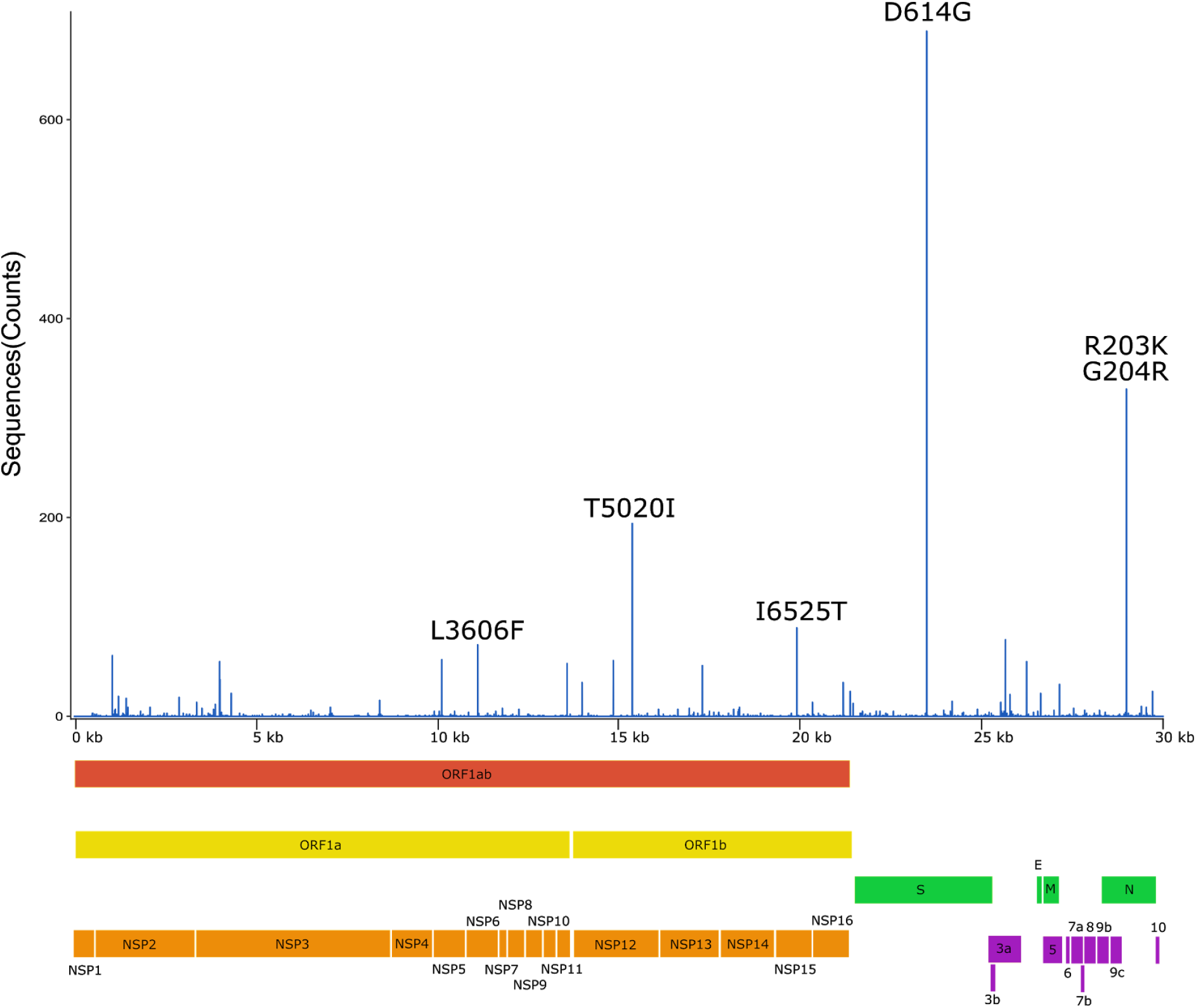
Graphical representation of the SARS-CoV-2 mutation prevalences in Belgium between February and mid June 2020. On top, a frequency plot displaying the quantity of sequences with a specific amino acid change (y-axis, counts). Amino acid changes found in at least 10% of the sequences (77, n= 776) are annotated. A scaled genetic map is displayed below, showing the different ORFs and protein products of SARS-CoV-2.

Prevalence of mutations L3606F, T5020I, I65256T, D614G, R203K and G204K varies over time (Figure 6). The amino acid substitution D614G, which is linked to an increased replicability of SARS-CoV-2 in Vero E6 cells^27^, was already prevalent in our sequencing space from the beginning of March (week 9). Moreover, the ratio of G614/D614 sequences increases over time, corresponding with data reported across Europe, where D614G was the most prevalent viral variant from April 2020 onwards (Figure 6) ^26^. As mentioned above, D614G is linked to a strong functional change, as the aspartic acid (D), an amino acid with bivalent cation binding properties, is exchanged for a glycine (G), a small apolar aminoacid, known to be flexible and capable of interrupting helicoidal secondary structures. This single non-synonymous mutation increases the fitness of the virus by increasing its replication in the upper respiratory tract. Increased titers in the upper respiratory tract coupled with a lack of increased morbidity makes D614G an adaptive variant for human-to-human transmission^14,27,28^. R203K and G204K are collinear and share the same evolutionary history in the Belgian set of sequences. K203/K204 are present in 31% of our sequences. Interestingly, both positions are found to change to a lysine (K). Unlike G204K, R203K corresponds to a conservative amino acid change. Both K and arginine (R) are basic amino acids with DNA/RNA binding properties. Both non-synonymous mutations are found between the RNA-Binding Domain (RBD) and the Dimerization Domain (BB), also called the link domain, of the nucleocapsid protein. Based on the high frequency of K203/K204 it appears that a local increase in positive polarity does not hinder N protein function. L3606F and T5020I (lineage B.1), causing amino acid changes in the NSP6 and NSP12 proteins, respectively, also seem to follow similar frequency trends over the first 20 weeks of the epidemic (Figure 6). Position 3606 of the ORF1ab (NSP6) gene undergoes a substantial change from a leucine (L), an aliphatic amino acid, to a phenylalanine (F), a more reactive aromatic amino acid. Position 5020 in the ORF1ab (NSP12) undergoes a strong change in hydrophobicity between a threonine (T), a polar hydroxylic amino acid and an isoleucine (I), an aliphatic hydrophobic amino acid (an increase in 0.493 in hydrophobicity). Similarly, the I6525T change in the ORF1ab (NSP15) results in a strong decrease in hydrophobicity. Currently, little is known about the functional consequences of these changes.

**Figure 6.**
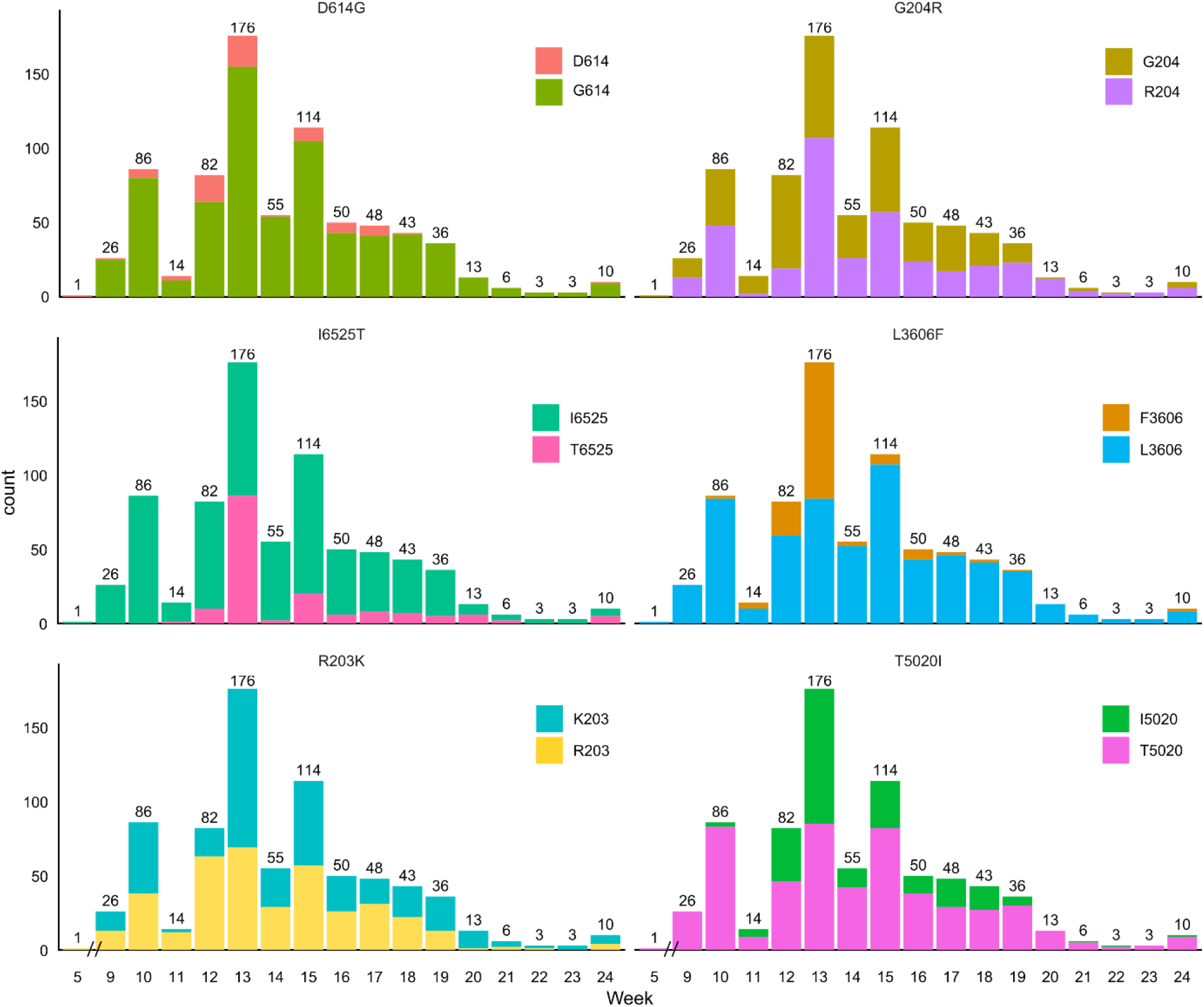
Temporal progression of the prevalence of the 6 most frequent mutations during the first pandemic wave of SARS-CoV-2 in Belgium. This scaled multipanel displays the number of sequences (y-axis, counts) divided between the 2 possible variants of each of the top 6 most frequent mutations during the first wave of SARS-CoV-2 in Belgium.

## Conclusion

The first wave of the SARS-CoV-2 epidemic in Belgium spanned from February 3rd (first reported case) to mid June. Over that period, 776 SARS-CoV-2 cases were sequenced, representing a randomized subset of the early circulation of SARS-CoV-2 in Belgium. Most reported COVID-19 cases corresponded to local or ‘domestic’ transmissions of SARS-CoV-2. Late March border control measures greatly contributed to reduce international import (i.e. returning travelers from holidays). Domestic clades and lineage distribution were comparable to the general European trend, with clades G, GR and GH being the most prominent. Despite the sequencing rate being relatively low during the peak of transmission of the first wave in Belgium (weeks 13 - 16, April 2020), it was sufficient to identify the presence of minor clades L, S and V, with S being mostly linked to direct SARS-CoV-2 import due to (i) its very low prevalence in Belgium (4 cases), (ii) its placement in a separate clade in the phylogenetic tree, as seen in Figure 3 and (iii) its direct link to travel history. Identification of 2 S clade sequences after border and flight regulations were already in place hints at the possibility of a small number of SARS-CoV-2-positive cases eluding the established measures. Furthermore, sequence resolution was sufficient to identify more than 42 SARS-CoV-2 lineages throughout the period, comparable to other reporting countries^26^. The most frequently observed non-conservative amino acid changes are in line with the presented clade distribution. Despite some of the mutations corresponding to significant changes in the chemical properties of the amino acids, little is known about their functional consequences on SARS-CoV-2 infection and COVID-19 presentation. Further efforts are required in (i) the detection of SARS-CoV-2 variants through random genomic surveillance to check measure effectiveness and viral evolution and in (ii) functional studies to map detected non-synonymous changes into viral behaviour.

## Acknowledgements

JMC is supported by a doctoral grant from HONOURs Marie-Sklodowska-Curie training network (721367). BV is supported by a FWO SB grant for strategic basic research of the Fonds Wetenschappelijk Onderzoek/Research Foundation Flanders (1S28617N). This work was supported by a COVID19 research grant of ‘Fonds Wetenschappelijk Onderzoek’/Research Foundation Flanders (G0H4420N). Part of this research was supported by “Internal Funds KU Leuven” awarded to PM (3M170314). UZ Leuven, as national reference center for respiratory pathogens, is supported by Sciensano, which is gratefully acknowledged. The authors acknowledge all research teams that have deposited SARS-CoV-2 genome data on GISAID (www.gisaid.org).

## Author contributions

TWB, JMC, BV, MB, PM, and EW conducted sample selection and processing. MB and EW conducted sample confirmation. LL, LC, KB and PM coordinated the collection and pre-selection of samples. TWB, JMC and BV conducted the sequencing. JMC and PM performed the data analysis and metadata curation. PM provided funding. JMC elaborated the manuscript draft. TWB, JMC, BV, MB, EW, LL, LC, KB, EA, MVR, and PM worked on the manuscript.

