## Supplementary_Material for "Genetic diversity and evolution of SARS-CoV-2 in Belgium during the first wave outbreak"

Suppl. Figure 1. Complete lineage classification of SARS-CoV-2 during the first pandemic wave in Belgium using the Pangolin tool (https://github.com/cov-lineages/pangolin).

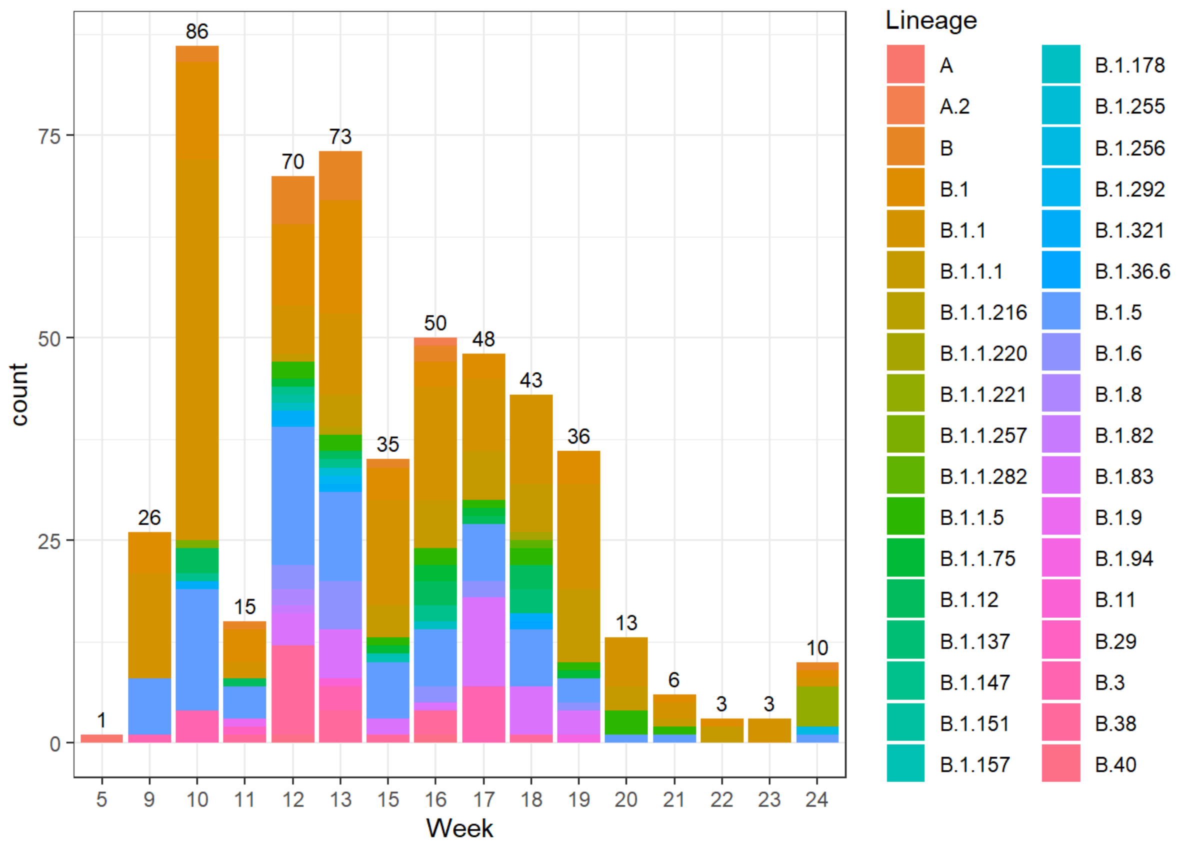

Suppl. Table 1. Collection of diagnosed and sequenced cases of SARS-CoV-2 in Belgium during its first epidemic wave. Data is distributed by weeks. SARS-CoV-2 positivity was collected from Sciensano public data resource (https://epistat.wiv-isp.be/covid/). Positive cases without an affiliated province have been excluded.

| Week | Cases | Sequenced |
|---|---|---|
| 5 | 1 | 1 |
| 9 | 72 | 26 |
| 10 | 510 | 86 |
| 11 | 1903 | 14 |
| 12 | 5297 | 82 |
| 13 | 8901 | 176 |
| 14 | 9527 | 55 |
| 15 | 9554 | 114 |
| 16 | 8220 | 50 |
| 17 | 4406 | 48 |
| 18 | 2927 | 43 |
| 19 | 2606 | 36 |
| 20 | 1748 | 13 |
| 21 | 1377 | 6 |
| 22 | 819 | 3 |
| 23 | 872 | 3 |
| 24 | 502 | 10 |

Suppl. Table 2. Lineage classification of SARS-CoV-2 genomes during the first infection wave in Belgium according to the Pangolin tool.

| SampleDate | Lineage | Month | Week |
|---|---|---|---|
| 03/02/2020 | A | Feb | 5 |
| 01/03/2020 | B.1.5 | Mar | 9 |
| 01/03/2020 | B.1.5 | Mar | 9 |
| 01/03/2020 | B.1.5 | Mar | 9 |
| 01/03/2020 | B.1.5 | Mar | 9 |
| 01/03/2020 | B.1.1 | Mar | 9 |
| 01/03/2020 | B.1.1 | Mar | 9 |
| 29/02/2020 | B.1 | Feb | 9 |
| 02/03/2020 | B.1.1 | Mar | 9 |
| 02/03/2020 | B.1.1 | Mar | 9 |
| 03/03/2020 | B.1 | Mar | 9 |
| 02/03/2020 | B.1 | Mar | 9 |
| 02/03/2020 | B.1.1 | Mar | 9 |
| 02/03/2020 | B.1.1 | Mar | 9 |
| 03/03/2020 | B.1.1 | Mar | 9 |
| 02/03/2020 | B.1.1 | Mar | 9 |
| 04/03/2020 | B.1.1 | Mar | 10 |
| 04/03/2020 | B.1.5 | Mar | 10 |
| 06/03/2020 | B.1.1 | Mar | 10 |
| 04/03/2020 | B.1.1 | Mar | 10 |
| 02/03/2020 | B.1.1 | Mar | 9 |
| 02/03/2020 | B.1.1 | Mar | 9 |
| 03/03/2020 | B.1.1 | Mar | 9 |
| 03/03/2020 | B.1 | Mar | 9 |
| 04/03/2020 | B.1.1 | Mar | 10 |
| 05/03/2020 | B.1 | Mar | 10 |
| 10/03/2020 | B.1 | Mar | 10 |
| 06/03/2020 | B.1.5 | Mar | 10 |
| 06/03/2020 | B.1.5 | Mar | 10 |
| 05/03/2020 | B.1.5 | Mar | 10 |
| 05/03/2020 | B.1.12 | Mar | 10 |
| 10/03/2020 | B.3 | Mar | 10 |
| 06/03/2020 | B | Mar | 10 |
| 05/03/2020 | B.1.5 | Mar | 10 |
| 05/03/2020 | B.1.1 | Mar | 10 |
| 06/03/2020 | B.1 | Mar | 10 |
| 09/03/2020 | B | Mar | 10 |
| 09/03/2020 | B.1.1.257 | Mar | 10 |
| 09/03/2020 | B.1.1 | Mar | 10 |
| 09/03/2020 | B.1.321 | Mar | 10 |
| 02/03/2020 | B.1.5 | Mar | 9 |

| | | | |
|---|---|---|---|
| 03/03/2020 | B.3 | Mar | 9 |
| 04/03/2020 | B.1.5 | Mar | 10 |
| 04/03/2020 | B.1 | Mar | 10 |
| 04/03/2020 | B.1.1 | Mar | 10 |
| 06/03/2020 | B.1.1 | Mar | 10 |
| 07/03/2020 | B.1.1 | Mar | 10 |
| 05/03/2020 | B.1.1 | Mar | 10 |
| 07/03/2020 | B.1.1 | Mar | 10 |
| 04/03/2020 | B.1.1 | Mar | 10 |
| 09/03/2020 | B.1.1 | Mar | 10 |
| 06/03/2020 | B.1.1 | Mar | 10 |
| 05/03/2020 | B.1 | Mar | 10 |
| 07/03/2020 | B.1.1 | Mar | 10 |
| 04/03/2020 | B.1 | Mar | 10 |
| 05/03/2020 | B.1 | Mar | 10 |
| 06/03/2020 | B.1.1 | Mar | 10 |
| 07/03/2020 | B.1 | Mar | 10 |
| 06/03/2020 | B.1.1 | Mar | 10 |
| 05/03/2020 | B.1.5 | Mar | 10 |
| 05/03/2020 | B.1.1 | Mar | 10 |
| 05/03/2020 | B.1.1 | Mar | 10 |
| 06/03/2020 | B.1.1 | Mar | 10 |
| 06/03/2020 | B.1.1 | Mar | 10 |
| 05/03/2020 | B.1.12 | Mar | 10 |
| 06/03/2020 | B.1.1 | Mar | 10 |
| 06/03/2020 | B.1.1 | Mar | 10 |
| 05/03/2020 | B.1.1 | Mar | 10 |
| 09/03/2020 | B.3 | Mar | 10 |
| 05/03/2020 | B.3 | Mar | 10 |
| 06/03/2020 | B.1.1 | Mar | 10 |
| 06/03/2020 | B.1.1 | Mar | 10 |
| 06/03/2020 | B.1 | Mar | 10 |
| 06/03/2020 | B.1.1 | Mar | 10 |
| 05/03/2020 | B.1.1 | Mar | 10 |
| 06/03/2020 | B.1.5 | Mar | 10 |
| 09/03/2020 | B.1.5 | Mar | 10 |
| 06/03/2020 | B.1 | Mar | 10 |
| 05/03/2020 | B.1.5 | Mar | 10 |
| 05/03/2020 | B.1.1 | Mar | 10 |
| 06/03/2020 | B.1.5 | Mar | 10 |
| 06/03/2020 | B.1.1 | Mar | 10 |
| 07/03/2020 | B.1.1 | Mar | 10 |
| 05/03/2020 | B.1.1 | Mar | 10 |
| 05/03/2020 | B.1.1 | Mar | 10 |
| 06/03/2020 | B.1.1 | Mar | 10 |
| 06/03/2020 | B.3 | Mar | 10 |

| | | | |
|---|---|---|---|
| 20/03/2020 | B.1.5 | Mar | 12 |
| 18/03/2020 | B.1 | Mar | 12 |
| 18/03/2020 | B.1 | Mar | 12 |
| 18/03/2020 | B.1 | Mar | 12 |
| 17/03/2020 | B.1 | Mar | 11 |
| 19/03/2020 | B.1 | Mar | 12 |
| 20/03/2020 | B.1.5 | Mar | 12 |
| 19/03/2020 | B.1.5 | Mar | 12 |
| 19/03/2020 | B.1.5 | Mar | 12 |
| 05/03/2020 | B.1.5 | Mar | 10 |
| 19/03/2020 | B.1.5 | Mar | 12 |
| 19/03/2020 | B.38 | Mar | 12 |
| 19/03/2020 | B.1.5 | Mar | 12 |
| 23/03/2020 | B | Mar | 12 |
| 23/03/2020 | B.1.151 | Mar | 12 |
| 23/03/2020 | B.1 | Mar | 12 |
| 24/03/2020 | B.40 | Mar | 12 |
| 24/03/2020 | B | Mar | 12 |
| 24/03/2020 | B.1.321 | Mar | 12 |
| 24/03/2020 | B.1.1.5 | Mar | 12 |
| 24/03/2020 | B | Mar | 12 |
| 25/03/2020 | B.1.1 | Mar | 13 |
| 25/03/2020 | B.1.83 | Mar | 13 |
| 24/03/2020 | B.1.5 | Mar | 12 |
| 24/03/2020 | B.1.83 | Mar | 12 |
| 24/03/2020 | B.1.321 | Mar | 12 |
| 25/03/2020 | B.1.5 | Mar | 13 |
| 25/03/2020 | B.1.6 | Mar | 13 |
| 23/03/2020 | B.1.5 | Mar | 12 |
| 24/03/2020 | B.38 | Mar | 12 |
| 24/03/2020 | B.1.5 | Mar | 12 |
| 24/03/2020 | B.1.83 | Mar | 12 |
| 24/03/2020 | B.1.5 | Mar | 12 |
| 24/03/2020 | B.1 | Mar | 12 |
| 24/03/2020 | B.38 | Mar | 12 |
| 24/03/2020 | B.1.1.1 | Mar | 12 |
| 25/03/2020 | B.1.6 | Mar | 13 |
| 24/03/2020 | B.1.82 | Mar | 12 |
| 23/03/2020 | B.1.83 | Mar | 12 |
| 23/03/2020 | B.1 | Mar | 12 |
| 23/03/2020 | B.1.6 | Mar | 12 |
| 24/03/2020 | B.1.1 | Mar | 12 |
| 25/03/2020 | B.1.1.1 | Mar | 13 |
| 24/03/2020 | B.1 | Mar | 12 |
| 25/03/2020 | B.1.1.1 | Mar | 13 |
| 25/03/2020 | B | Mar | 13 |

| | | | |
|---|---|---|---|
| 25/03/2020 | B.1.1 | Mar | 13 |
| 25/03/2020 | B.1.1.5 | Mar | 13 |
| 24/03/2020 | B.1.83 | Mar | 12 |
| 24/03/2020 | B.1.6 | Mar | 12 |
| 24/03/2020 | B.1.1.1 | Mar | 12 |
| 25/03/2020 | B.1.147 | Mar | 13 |
| 25/03/2020 | B.1.1 | Mar | 13 |
| 23/03/2020 | B.1.1 | Mar | 12 |
| 24/03/2020 | B | Mar | 12 |
| 25/03/2020 | B.1.292 | Mar | 13 |
| 25/03/2020 | B.1 | Mar | 13 |
| 25/03/2020 | B.1.6 | Mar | 13 |
| 23/03/2020 | B.1.1 | Mar | 12 |
| 24/03/2020 | B.1.12 | Mar | 12 |
| 24/03/2020 | B.1.1.5 | Mar | 12 |
| 25/03/2020 | B.1 | Mar | 13 |
| 24/03/2020 | B.1 | Mar | 12 |
| 24/03/2020 | B.1.1.5 | Mar | 12 |
| 25/03/2020 | B | Mar | 13 |
| 25/03/2020 | B.1 | Mar | 13 |
| 24/03/2020 | B.1.83 | Mar | 12 |
| 24/03/2020 | B.1.1 | Mar | 12 |
| 24/03/2020 | B.1.1 | Mar | 12 |
| 24/03/2020 | B.1.5 | Mar | 12 |
| 25/03/2020 | B.1.321 | Mar | 13 |
| 24/03/2020 | B.1.6 | Mar | 12 |
| 24/03/2020 | B.1 | Mar | 12 |
| 24/03/2020 | B.1 | Mar | 12 |
| 23/03/2020 | B.1.5 | Mar | 12 |
| 24/03/2020 | B.1.1 | Mar | 12 |
| 24/03/2020 | B.1 | Mar | 12 |
| 25/03/2020 | B.1 | Mar | 13 |
| 24/03/2020 | B.1.6 | Mar | 12 |
| 25/03/2020 | B | Mar | 13 |
| 25/03/2020 | B | Mar | 13 |
| 25/03/2020 | B.1.1 | Mar | 13 |
| 25/03/2020 | B.1.5 | Mar | 13 |
| 06/03/2020 | B.1.1 | Mar | 10 |
| 04/03/2020 | B.1.12 | Mar | 10 |
| 04/03/2020 | B.1.5 | Mar | 10 |
| 05/03/2020 | B.1.1 | Mar | 10 |
| 09/03/2020 | B.1.147 | Mar | 10 |
| 06/03/2020 | B.1.1 | Mar | 10 |
| 04/03/2020 | B.1.1 | Mar | 10 |
| 04/03/2020 | B.1.1 | Mar | 10 |
| 06/03/2020 | B.1 | Mar | 10 |

| | | | |
|---|---|---|---|
| 07/03/2020 | B.1.1 | Mar | 10 |
| 07/03/2020 | B.1.1 | Mar | 10 |
| 09/03/2020 | B.1.5 | Mar | 10 |
| 05/03/2020 | B.1.1 | Mar | 10 |
| 04/03/2020 | B.1.1 | Mar | 10 |
| 28/03/2020 | B.1.1.216 | Mar | 13 |
| 28/03/2020 | B.1.1 | Mar | 13 |
| 28/03/2020 | B.1.1 | Mar | 13 |
| 28/03/2020 | B.1.5 | Mar | 13 |
| 28/03/2020 | B.1.83 | Mar | 13 |
| 28/03/2020 | B.1.6 | Mar | 13 |
| 28/03/2020 | B.1.83 | Mar | 13 |
| 28/03/2020 | B.1.83 | Mar | 13 |
| 28/03/2020 | B.1.5 | Mar | 13 |
| 28/03/2020 | B.1.5 | Mar | 13 |
| 28/03/2020 | B.1 | Mar | 13 |
| 28/03/2020 | B.1.83 | Mar | 13 |
| 28/03/2020 | B.1.1 | Mar | 13 |
| 28/03/2020 | B.1.83 | Mar | 13 |
| 28/03/2020 | B.1.1.75 | Mar | 13 |
| 28/03/2020 | B.1.1 | Mar | 13 |
| 28/03/2020 | B.1.83 | Mar | 13 |
| 28/03/2020 | B.1 | Mar | 13 |
| 28/03/2020 | B.1 | Mar | 13 |
| 28/03/2020 | B.1.6 | Mar | 13 |
| 20/03/2020 | B.1.5 | Mar | 12 |
| 28/03/2020 | B.1 | Mar | 13 |
| 28/03/2020 | B.1.1.1 | Mar | 13 |
| 28/03/2020 | B.1.178 | Mar | 13 |
| 27/03/2020 | B.1.1.1 | Mar | 13 |
| 27/03/2020 | B.1 | Mar | 13 |
| 27/03/2020 | B.1.6 | Mar | 13 |
| 27/03/2020 | B.1.5 | Mar | 13 |
| 27/03/2020 | B.1 | Mar | 13 |
| 27/03/2020 | B.1.1.1 | Mar | 13 |
| 27/03/2020 | B.1.83 | Mar | 13 |
| 29/03/2020 | B.1.1 | Mar | 13 |
| 29/03/2020 | B.1.321 | Mar | 13 |
| 29/03/2020 | B.1.5 | Mar | 13 |
| 29/03/2020 | B.1.83 | Mar | 13 |
| 29/03/2020 | B.1 | Mar | 13 |
| 29/03/2020 | B.1.83 | Mar | 13 |
| 29/03/2020 | B.1.12 | Mar | 13 |
| 29/03/2020 | B.1.12 | Mar | 13 |
| 29/03/2020 | B.1 | Mar | 13 |
| 29/03/2020 | B.1.83 | Mar | 13 |

| | | | |
|---|---|---|---|
| 29/03/2020 | B.1.83 | Mar | 13 |
| 29/03/2020 | B.1.178 | Mar | 13 |
| 29/03/2020 | B.1.83 | Mar | 13 |
| 29/03/2020 | B.1 | Mar | 13 |
| 29/03/2020 | B.1 | Mar | 13 |
| 29/03/2020 | B | Mar | 13 |
| 29/03/2020 | B.1.1.5 | Mar | 13 |
| 29/03/2020 | B | Mar | 13 |
| 29/03/2020 | B.1.83 | Mar | 13 |
| 29/03/2020 | B.1.5 | Mar | 13 |
| 29/03/2020 | B.1.5 | Mar | 13 |
| 29/03/2020 | B.1.1 | Mar | 13 |
| 29/03/2020 | B.1.5 | Mar | 13 |
| 29/03/2020 | B | Mar | 13 |
| 29/03/2020 | B.1.1 | Mar | 13 |
| 29/03/2020 | B.1.1 | Mar | 13 |
| 29/03/2020 | B.1 | Mar | 13 |
| 29/03/2020 | B.1.12 | Mar | 13 |
| 29/03/2020 | B.1 | Mar | 13 |
| 29/03/2020 | B.1 | Mar | 13 |
| 30/03/2020 | B.1.1.75 | Mar | 13 |
| 30/03/2020 | B.1.83 | Mar | 13 |
| 30/03/2020 | B.1.1.5 | Mar | 13 |
| 30/03/2020 | B.1 | Mar | 13 |
| 30/03/2020 | B | Mar | 13 |
| 30/03/2020 | B.1.1.1 | Mar | 13 |
| 30/03/2020 | B.1.83 | Mar | 13 |
| 30/03/2020 | B.1.1.75 | Mar | 13 |
| 30/03/2020 | B.3 | Mar | 13 |
| 30/03/2020 | B.1.1 | Mar | 13 |
| 30/03/2020 | B.1.1.75 | Mar | 13 |
| 20/03/2020 | B.1.147 | Mar | 12 |
| 19/03/2020 | B.1.1.5 | Mar | 12 |
| 16/03/2020 | B.1.1 | Mar | 11 |
| 26/03/2020 | B.1 | Mar | 13 |
| 20/03/2020 | B | Mar | 12 |
| 31/03/2020 | B.1 | Mar | 13 |
| 31/03/2020 | B.1.1.5 | Mar | 13 |
| 19/03/2020 | B.1 | Mar | 12 |
| 23/04/2020 | B.3 | Apr | 17 |
| 23/04/2020 | B.3 | Apr | 17 |
| 23/04/2020 | B.3 | Apr | 17 |
| 23/04/2020 | B.3 | Apr | 17 |
| 23/04/2020 | B.3 | Apr | 17 |
| 23/04/2020 | B.3 | Apr | 17 |
| 23/04/2020 | B.3 | Apr | 17 |

| | | | |
|---|---|---|---|
| 30/03/2020 | B.1 | Mar | 13 |
| 30/03/2020 | B | Mar | 13 |
| 30/03/2020 | B.1 | Mar | 13 |
| 30/03/2020 | B.1.8 | Mar | 13 |
| 20/03/2020 | B.1.1 | Mar | 12 |
| 28/03/2020 | B.1.5 | Mar | 13 |
| 28/03/2020 | B.1 | Mar | 13 |
| 28/03/2020 | B.1 | Mar | 13 |
| 28/03/2020 | B.1.13 | Mar | 13 |
| 28/03/2020 | B.1.5 | Mar | 13 |
| 28/03/2020 | B.1.5 | Mar | 13 |
| 26/03/2020 | B.1 | Mar | 13 |
| 26/03/2020 | B.1.5 | Mar | 13 |
| 27/03/2020 | B.1.5 | Mar | 13 |
| 27/03/2020 | B.1.255 | Mar | 13 |
| 27/03/2020 | B.1 | Mar | 13 |
| 27/03/2020 | B.1 | Mar | 13 |
| 27/03/2020 | B.1.5 | Mar | 13 |
| 26/03/2020 | B.38 | Mar | 13 |
| 26/03/2020 | B.1 | Mar | 13 |
| 17/03/2020 | B.1.5 | Mar | 11 |
| 17/03/2020 | B.1.12 | Mar | 11 |
| 17/03/2020 | B.1.5 | Mar | 11 |
| 30/03/2020 | B | Mar | 13 |
| 30/03/2020 | B | Mar | 13 |
| 30/03/2020 | B.1.1 | Mar | 13 |
| 30/03/2020 | B | Mar | 13 |
| 30/03/2020 | B | Mar | 13 |
| 24/03/2020 | B.1.1.75 | Mar | 12 |
| 25/03/2020 | B.1 | Mar | 13 |
| 30/03/2020 | B.1.12 | Mar | 13 |
| 26/03/2020 | B.3 | Mar | 13 |
| 25/03/2020 | B.1.5 | Mar | 13 |
| 28/03/2020 | B.1.22 | Mar | 13 |
| 25/03/2020 | B.1.1 | Mar | 13 |
| 25/03/2020 | B | Mar | 13 |
| 28/03/2020 | B.1.1 | Mar | 13 |
| 28/03/2020 | B.1.1 | Mar | 13 |
| 28/03/2020 | B.1.1 | Mar | 13 |
| 27/03/2020 | B.1.1.5 | Mar | 13 |
| 27/03/2020 | B.1.1 | Mar | 13 |
| 27/03/2020 | B.1.1 | Mar | 13 |
| 30/03/2020 | B.1.1 | Mar | 13 |
| 27/03/2020 | B | Mar | 13 |
| 27/03/2020 | B.11 | Mar | 13 |
| 26/03/2020 | B.1.5 | Mar | 13 |

| | | | |
|---|---|---|---|
| 30/03/2020 | B.1.5 | Mar | 13 |
| 29/03/2020 | B.1.1 | Mar | 13 |
| 27/03/2020 | B.1.12 | Mar | 13 |
| 28/03/2020 | B.1 | Mar | 13 |
| 22/03/2020 | B | Mar | 12 |
| 11/03/2020 | B.1 | Mar | 11 |
| 29/03/2020 | B.1.1.74 | Mar | 13 |
| 30/03/2020 | B.1.1 | Mar | 13 |
| 31/03/2020 | B.1.1.1 | Mar | 13 |
| 31/03/2020 | B.1.147 | Mar | 13 |
| 31/03/2020 | B.38 | Mar | 13 |
| 31/03/2020 | B.1.5 | Mar | 13 |
| 30/03/2020 | B.1.9 | Mar | 13 |
| 01/04/2020 | B.1.147 | Apr | 14 |
| 30/03/2020 | B.1.12 | Mar | 13 |
| 01/04/2020 | B.1 | Apr | 14 |
| 30/03/2020 | B.1.9 | Mar | 13 |
| 01/04/2020 | B.1.1 | Apr | 14 |
| 27/03/2020 | B.3 | Mar | 13 |
| 28/03/2020 | B.1.1 | Mar | 13 |
| 01/04/2020 | B.1 | Apr | 14 |
| 01/04/2020 | B.1.5 | Apr | 14 |
| 01/04/2020 | B.1 | Apr | 14 |
| 02/04/2020 | B.1.1 | Apr | 14 |
| 01/04/2020 | B.1.1 | Apr | 14 |
| 01/04/2020 | B.1.1 | Apr | 14 |
| 31/03/2020 | B.1.1 | Mar | 13 |
| 31/03/2020 | B.1.353 | Mar | 13 |
| 31/03/2020 | B.1.5 | Mar | 13 |
| 25/03/2020 | B.1.6 | Mar | 13 |
| 03/04/2020 | B.1.5 | Apr | 14 |
| 03/04/2020 | B.1.5 | Apr | 14 |
| 02/04/2020 | B.1.6 | Apr | 14 |
| 30/03/2020 | B.1.1 | Mar | 13 |
| 30/03/2020 | B.1.1.189 | Mar | 13 |
| 29/03/2020 | B.1.1.75 | Mar | 13 |
| 30/03/2020 | B.1.1.75 | Mar | 13 |
| 28/03/2020 | B | Mar | 13 |
| 29/03/2020 | B.1.1 | Mar | 13 |
| 27/03/2020 | B.3 | Mar | 13 |
| 26/03/2020 | B.1 | Mar | 13 |
| 29/03/2020 | B.1.1 | Mar | 13 |
| 07/04/2020 | B.1.1.1 | Apr | 14 |
| 07/04/2020 | B.1 | Apr | 14 |
| 07/04/2020 | B.1.1.5 | Apr | 14 |
| 07/04/2020 | B.1.1.1 | Apr | 14 |

| | | | |
|---|---|---|---|
| 07/04/2020 | B.1.1.1 | Apr | 14 |
| 06/04/2020 | B.1.5 | Apr | 14 |
| 08/04/2020 | B.1 | Apr | 15 |
| 06/04/2020 | B.1.5 | Apr | 14 |
| 12/03/2020 | B.1 | Mar | 11 |
| 25/03/2020 | B.1.1 | Mar | 13 |
| 07/04/2020 | B.1.5 | Apr | 14 |
| 07/04/2020 | B.1 | Apr | 14 |
| 07/04/2020 | B.1 | Apr | 14 |
| 09/04/2020 | B.1.83 | Apr | 15 |
| 09/04/2020 | B.1.83 | Apr | 15 |
| 11/04/2020 | B.1.5 | Apr | 15 |
| 23/03/2020 | B.1.8 | Mar | 12 |
| 23/03/2020 | B | Mar | 12 |
| 22/03/2020 | B.1.8 | Mar | 12 |
| 08/04/2020 | B.1.147 | Apr | 15 |
| 08/04/2020 | B.1.147 | Apr | 15 |
| 08/04/2020 | B.1.5 | Apr | 15 |
| 09/04/2020 | B.38 | Apr | 15 |
| 06/05/2020 | B.1.1 | May | 19 |
| 07/05/2020 | B.1.1 | May | 19 |
| 07/05/2020 | B.1.1.1 | May | 19 |
| 07/05/2020 | B.1.1.1 | May | 19 |
| 07/05/2020 | B.1.1.1 | May | 19 |
| 07/05/2020 | B.1.5 | May | 19 |
| 07/04/2020 | B.1.1.221 | Apr | 14 |
| 07/04/2020 | B.1.1.1 | Apr | 14 |
| 06/04/2020 | B.1 | Apr | 14 |
| 07/04/2020 | B.1.1 | Apr | 14 |
| 06/04/2020 | B.1.1 | Apr | 14 |
| 04/04/2020 | B.1 | Apr | 14 |
| 04/04/2020 | B.1.12 | Apr | 14 |
| 06/04/2020 | B.1.83 | Apr | 14 |
| 06/04/2020 | B.1.1.1 | Apr | 14 |
| 07/04/2020 | B.1 | Apr | 14 |
| 08/04/2020 | B.3 | Apr | 15 |
| 07/04/2020 | B.1.6 | Apr | 14 |
| 07/04/2020 | B.1.83 | Apr | 14 |
| 08/04/2020 | B.1.1 | Apr | 15 |
| 06/04/2020 | B.38 | Apr | 14 |
| 08/04/2020 | B.1.1.1 | Apr | 15 |
| 07/04/2020 | B.1 | Apr | 14 |
| 07/04/2020 | B.1.5 | Apr | 14 |
| 05/04/2020 | B.1.321 | Apr | 14 |
| 03/04/2020 | B.1 | Apr | 14 |
| 09/04/2020 | B.1.1 | Apr | 15 |

| | | | |
|---|---|---|---|
| 08/04/2020 | B.1.83 | Apr | 15 |
| 08/04/2020 | B.1.83 | Apr | 15 |
| 09/04/2020 | B.1.1 | Apr | 15 |
| 09/04/2020 | B | Apr | 15 |
| 09/04/2020 | B.1.5 | Apr | 15 |
| 09/04/2020 | B.1.1 | Apr | 15 |
| 09/04/2020 | B.1.1 | Apr | 15 |
| 09/04/2020 | B.1.1 | Apr | 15 |
| 09/04/2020 | B.1.5 | Apr | 15 |
| 09/04/2020 | B.1.1 | Apr | 15 |
| 09/04/2020 | B.1.1.1 | Apr | 15 |
| 09/04/2020 | B.1.1.5 | Apr | 15 |
| 09/04/2020 | B.1.1 | Apr | 15 |
| 09/04/2020 | B.1.1.75 | Apr | 15 |
| 11/04/2020 | B.1.1 | Apr | 15 |
| 11/04/2020 | B.1.5 | Apr | 15 |
| 11/04/2020 | B.1.5 | Apr | 15 |
| 11/04/2020 | B.40 | Apr | 15 |
| 11/04/2020 | B.1.1.221 | Apr | 15 |
| 11/04/2020 | B.1.1.1 | Apr | 15 |
| 11/04/2020 | B.1.83 | Apr | 15 |
| 11/04/2020 | B.1.1 | Apr | 15 |
| 11/04/2020 | B.1.1.1 | Apr | 15 |
| 11/04/2020 | B.1.1 | Apr | 15 |
| 11/04/2020 | B.1.147 | Apr | 15 |
| 11/04/2020 | B.1.147 | Apr | 15 |
| 11/04/2020 | B | Apr | 15 |
| 11/04/2020 | B.1.321 | Apr | 15 |
| 11/04/2020 | B.1.1 | Apr | 15 |
| 11/04/2020 | B.1.147 | Apr | 15 |
| 11/04/2020 | B.1.1.221 | Apr | 15 |
| 11/04/2020 | B.1.12 | Apr | 15 |
| 11/04/2020 | B.1.292 | Apr | 15 |
| 11/04/2020 | B.1.1.5 | Apr | 15 |
| 11/04/2020 | B.1.1.75 | Apr | 15 |
| 11/04/2020 | B.1.321 | Apr | 15 |
| 11/04/2020 | B.1.321 | Apr | 15 |
| 11/04/2020 | B.1.1.1 | Apr | 15 |
| 11/04/2020 | B.1.6 | Apr | 15 |
| 11/04/2020 | B.1.6 | Apr | 15 |
| 11/04/2020 | B.1.1 | Apr | 15 |
| 11/04/2020 | B.1 | Apr | 15 |
| 11/04/2020 | B.1.1.1 | Apr | 15 |
| 11/04/2020 | B.1.1 | Apr | 15 |
| 11/04/2020 | B.1.5 | Apr | 15 |
| 11/04/2020 | B.1.1 | Apr | 15 |

| | | | |
|---|---|---|---|
| 11/04/2020 | B.1.1 | Apr | 15 |
| 14/04/2020 | B.1.1.75 | Apr | 15 |
| 14/04/2020 | B.1.1 | Apr | 15 |
| 14/04/2020 | B | Apr | 15 |
| 14/04/2020 | B.1.1.1 | Apr | 15 |
| 14/04/2020 | B.1.1.1 | Apr | 15 |
| 14/04/2020 | B.1.5 | Apr | 15 |
| 14/04/2020 | B.1.5 | Apr | 15 |
| 18/04/2020 | B.1.1.1 | Apr | 16 |
| 21/04/2020 | B.1.5 | Apr | 16 |
| 27/04/2020 | B.1.1.1 | Apr | 17 |
| 28/04/2020 | B.1 | Apr | 17 |
| 28/04/2020 | B.1.83 | Apr | 17 |
| 29/04/2020 | B.1.83 | Apr | 18 |
| 01/05/2020 | B.1.1.1 | May | 18 |
| 02/05/2020 | B.1.5 | May | 18 |
| 02/05/2020 | B.1.1 | May | 18 |
| 03/05/2020 | B.1.5 | May | 18 |
| 04/05/2020 | B.1.1 | May | 18 |
| 05/05/2020 | B.1.1.220 | May | 18 |
| 05/05/2020 | B.1.1.5 | May | 18 |
| 06/05/2020 | B.1.1 | May | 19 |
| 07/05/2020 | B.1.1 | May | 19 |
| 11/05/2020 | B.1 | May | 19 |
| 13/05/2020 | B.1.1.5 | May | 20 |
| 09/03/2020 | B.1.1 | Mar | 10 |
| 10/06/2020 | B | Jun | 24 |
| 11/03/2020 | B.1.1 | Mar | 11 |
| 29/03/2020 | B.1 | Mar | 13 |
| 03/03/2020 | B.1.1 | Mar | 9 |
| 11/05/2020 | B.1.5 | May | 19 |
| 11/06/2020 | B.1.5 | Jun | 24 |
| 29/04/2020 | B.1.5 | Apr | 18 |
| 03/03/2020 | B.1.5 | Mar | 9 |
| 12/03/2020 | B.1.5 | Mar | 11 |
| 15/03/2020 | B.1.5 | Mar | 11 |
| 17/03/2020 | B.38 | Mar | 11 |
| 19/03/2020 | B.38 | Mar | 12 |
| 19/03/2020 | B.38 | Mar | 12 |
| 19/03/2020 | B.1.5 | Mar | 12 |
| 20/03/2020 | B.1.5 | Mar | 12 |
| 20/03/2020 | B.38 | Mar | 12 |
| 20/03/2020 | B.38 | Mar | 12 |
| 20/03/2020 | B.38 | Mar | 12 |
| 22/03/2020 | B.38 | Mar | 12 |
| 22/03/2020 | B.38 | Mar | 12 |

| | | | |
|---|---|---|---|
| 22/03/2020 | B.38 | Mar | 12 |
| 23/03/2020 | B.1.5 | Mar | 12 |
| 23/03/2020 | B.1.5 | Mar | 12 |
| 23/03/2020 | B.1.5 | Mar | 12 |
| 24/03/2020 | B.1.1 | Mar | 12 |
| 25/03/2020 | B.38 | Mar | 13 |
| 25/03/2020 | B.38 | Mar | 13 |
| 26/03/2020 | B.38 | Mar | 13 |
| 28/03/2020 | B.1.1.285 | Mar | 13 |
| 30/03/2020 | B.1.1 | Mar | 13 |
| 31/03/2020 | B.1.1 | Mar | 13 |
| 31/03/2020 | B.1.1 | Mar | 13 |
| 01/04/2020 | B.1.1 | Apr | 14 |
| 01/04/2020 | B.1.5 | Apr | 14 |
| 01/04/2020 | B.1.1 | Apr | 14 |
| 02/04/2020 | B.1.1 | Apr | 14 |
| 03/04/2020 | B.1.1 | Apr | 14 |
| 03/04/2020 | B.1.1 | Apr | 14 |
| 03/04/2020 | B.1.1 | Apr | 14 |
| 04/04/2020 | B.1.1 | Apr | 14 |
| 06/04/2020 | B.1.5 | Apr | 14 |
| 06/04/2020 | B.1 | Apr | 14 |
| 06/04/2020 | B.1.1 | Apr | 14 |
| 06/04/2020 | B.1.1 | Apr | 14 |
| 07/04/2020 | B.1.1 | Apr | 14 |
| 07/04/2020 | B.1.1 | Apr | 14 |
| 07/04/2020 | B.1.1 | Apr | 14 |
| 07/04/2020 | B.1.1 | Apr | 14 |
| 08/04/2020 | B.1.1 | Apr | 15 |
| 08/04/2020 | B.1.1 | Apr | 15 |
| 08/04/2020 | B.1.1 | Apr | 15 |
| 08/04/2020 | B.1.1 | Apr | 15 |
| 08/04/2020 | B.1.1 | Apr | 15 |
| 08/04/2020 | B.1.1 | Apr | 15 |
| 09/04/2020 | B.1.5 | Apr | 15 |
| 09/04/2020 | B.1.5 | Apr | 15 |
| 09/04/2020 | B.1.5 | Apr | 15 |
| 09/04/2020 | B.3 | Apr | 15 |
| 09/04/2020 | B.1.5 | Apr | 15 |
| 09/04/2020 | B.1.1 | Apr | 15 |
| 09/04/2020 | B.1.1.5 | Apr | 15 |
| 09/04/2020 | B.1.1 | Apr | 15 |
| 09/04/2020 | B.1.1 | Apr | 15 |
| 09/04/2020 | B.1.6 | Apr | 15 |
| 10/04/2020 | B.1 | Apr | 15 |
| 10/04/2020 | B.1.12 | Apr | 15 |

| | | | |
|---|---|---|---|
| 10/04/2020 | B.1.5 | Apr | 15 |
| 10/04/2020 | B.1.1 | Apr | 15 |
| 10/04/2020 | B.1.1 | Apr | 15 |
| 11/04/2020 | B.38 | Apr | 15 |
| 11/04/2020 | B.1.1 | Apr | 15 |
| 12/04/2020 | B.1.1 | Apr | 15 |
| 13/04/2020 | B.1 | Apr | 15 |
| 13/04/2020 | B.1.1 | Apr | 15 |
| 13/04/2020 | B.1.1 | Apr | 15 |
| 13/04/2020 | B.1 | Apr | 15 |
| 14/04/2020 | B.1.5 | Apr | 15 |
| 14/04/2020 | B.1 | Apr | 15 |
| 14/04/2020 | B.1 | Apr | 15 |
| 14/04/2020 | B.1.1 | Apr | 15 |
| 14/04/2020 | B.1.83 | Apr | 15 |
| 14/04/2020 | B.1.5 | Apr | 15 |
| 14/04/2020 | B.1.1 | Apr | 15 |
| 14/04/2020 | B.1.83 | Apr | 15 |
| 14/04/2020 | B.1.1.1 | Apr | 15 |
| 14/04/2020 | B.1.1.1 | Apr | 15 |
| 14/04/2020 | B.1.1 | Apr | 15 |
| 14/04/2020 | B.1.1 | Apr | 15 |
| 14/04/2020 | B.1.1 | Apr | 15 |
| 14/04/2020 | B.1.1 | Apr | 15 |
| 15/04/2020 | B.1.1 | Apr | 16 |
| 15/04/2020 | B.1.1 | Apr | 16 |
| 16/04/2020 | B.1 | Apr | 16 |
| 16/04/2020 | B.1.12 | Apr | 16 |
| 16/04/2020 | B.1.5 | Apr | 16 |
| 16/04/2020 | A.2 | Apr | 16 |
| 16/04/2020 | B.1.5 | Apr | 16 |
| 16/04/2020 | B.40 | Apr | 16 |
| 16/04/2020 | B.1.1.75 | Apr | 16 |
| 16/04/2020 | B.38 | Apr | 16 |
| 17/04/2020 | B.1.1 | Apr | 16 |
| 17/04/2020 | B.1.5 | Apr | 16 |
| 17/04/2020 | B.1.1 | Apr | 16 |
| 17/04/2020 | B.1.1 | Apr | 16 |
| 17/04/2020 | B.1.6 | Apr | 16 |
| 17/04/2020 | B.1.5 | Apr | 16 |
| 17/04/2020 | B.1.1 | Apr | 16 |
| 17/04/2020 | B.1.1 | Apr | 16 |
| 17/04/2020 | B.1.1.75 | Apr | 16 |
| 17/04/2020 | B.1 | Apr | 16 |
| 17/04/2020 | B.1.6 | Apr | 16 |
| 17/04/2020 | B.1.1.5 | Apr | 16 |

| | | | |
|---|---|---|---|
| 17/04/2020 | B.1.12 | Apr | 16 |
| 17/04/2020 | B.1.5 | Apr | 16 |
| 17/04/2020 | B | Apr | 16 |
| 17/04/2020 | B.38 | Apr | 16 |
| 17/04/2020 | B.1.147 | Apr | 16 |
| 17/04/2020 | B.1.147 | Apr | 16 |
| 17/04/2020 | B.1.1.1 | Apr | 16 |
| 17/04/2020 | B.1.1 | Apr | 16 |
| 17/04/2020 | B.1.1 | Apr | 16 |
| 17/04/2020 | B | Apr | 16 |
| 17/04/2020 | B.1.1 | Apr | 16 |
| 17/04/2020 | B.1.1 | Apr | 16 |
| 18/04/2020 | B.1.12 | Apr | 16 |
| 18/04/2020 | B.1.178 | Apr | 16 |
| 18/04/2020 | B.1.1.1 | Apr | 16 |
| 19/04/2020 | B.1.1.1 | Apr | 16 |
| 19/04/2020 | B.1.1 | Apr | 16 |
| 19/04/2020 | B.38 | Apr | 16 |
| 19/04/2020 | B.1.1 | Apr | 16 |
| 19/04/2020 | B.1.83 | Apr | 16 |
| 19/04/2020 | B.1.1.1 | Apr | 16 |
| 19/04/2020 | B.1.1.5 | Apr | 16 |
| 19/04/2020 | B.1.1 | Apr | 16 |
| 19/04/2020 | B.1.5 | Apr | 16 |
| 20/04/2020 | B.1 | Apr | 16 |
| 22/04/2020 | B.1.6 | Apr | 17 |
| 22/04/2020 | B.1.1 | Apr | 17 |
| 22/04/2020 | B.1 | Apr | 17 |
| 22/04/2020 | B.1.1.5 | Apr | 17 |
| 22/04/2020 | B.1.1 | Apr | 17 |
| 22/04/2020 | B.1.1.1 | Apr | 17 |
| 22/04/2020 | B.1.5 | Apr | 17 |
| 23/04/2020 | B.1.1 | Apr | 17 |
| 23/04/2020 | B.1.1.75 | Apr | 17 |
| 23/04/2020 | B.1.6 | Apr | 17 |
| 23/04/2020 | B.1 | Apr | 17 |
| 23/04/2020 | B.1.5 | Apr | 17 |
| 23/04/2020 | B.1.1 | Apr | 17 |
| 23/04/2020 | B.1.5 | Apr | 17 |
| 23/04/2020 | B.1.1 | Apr | 17 |
| 23/04/2020 | B.1.1.1 | Apr | 17 |
| 24/04/2020 | B.1.1.1 | Apr | 17 |
| 26/04/2020 | B.1.83 | Apr | 17 |
| 26/04/2020 | B.1.12 | Apr | 17 |
| 26/04/2020 | B.1.1.1 | Apr | 17 |
| 26/04/2020 | B.1.83 | Apr | 17 |

| | | | |
|---|---|---|---|
| 26/04/2020 | B.1.5 | Apr | 17 |
| 26/04/2020 | B.1.83 | Apr | 17 |
| 26/04/2020 | B.1.1 | Apr | 17 |
| 27/04/2020 | B.1.83 | Apr | 17 |
| 27/04/2020 | B.1.1 | Apr | 17 |
| 27/04/2020 | B.1.83 | Apr | 17 |
| 27/04/2020 | B.1.1 | Apr | 17 |
| 27/04/2020 | B.1.5 | Apr | 17 |
| 27/04/2020 | B.1.5 | Apr | 17 |
| 27/04/2020 | B.1.1 | Apr | 17 |
| 27/04/2020 | B.1.83 | Apr | 17 |
| 27/04/2020 | B.1.83 | Apr | 17 |
| 27/04/2020 | B.1.83 | Apr | 17 |
| 27/04/2020 | B.1.83 | Apr | 17 |
| 27/04/2020 | B.1.1.1 | Apr | 17 |
| 27/04/2020 | B.1.5 | Apr | 17 |
| 28/04/2020 | B.1.83 | Apr | 17 |
| 29/04/2020 | B.1.5 | Apr | 18 |
| 29/04/2020 | B.38 | Apr | 18 |
| 29/04/2020 | B.1.12 | Apr | 18 |
| 29/04/2020 | B.1.12 | Apr | 18 |
| 29/04/2020 | B.1.1 | Apr | 18 |
| 29/04/2020 | B.1.1 | Apr | 18 |
| 29/04/2020 | B.1.137 | Apr | 18 |
| 29/04/2020 | B.1.1 | Apr | 18 |
| 29/04/2020 | B.1.83 | Apr | 18 |
| 29/04/2020 | B.1.1 | Apr | 18 |
| 29/04/2020 | B.1.137 | Apr | 18 |
| 29/04/2020 | B.1.1.1 | Apr | 18 |
| 29/04/2020 | B.1.137 | Apr | 18 |
| 29/04/2020 | B.1.1.282 | Apr | 18 |
| 29/04/2020 | B.1.1 | Apr | 18 |
| 29/04/2020 | B.1.5 | Apr | 18 |
| 30/04/2020 | B.1.1 | Apr | 18 |
| 30/04/2020 | B.1.1.1 | Apr | 18 |
| 01/05/2020 | B.1.36.6 | May | 18 |
| 01/05/2020 | B.1.12 | May | 18 |
| 01/05/2020 | B.1.5 | May | 18 |
| 01/05/2020 | B.1.83 | May | 18 |
| 01/05/2020 | B.1.321 | May | 18 |
| 01/05/2020 | B.1.1 | May | 18 |
| 02/05/2020 | B.1.83 | May | 18 |
| 02/05/2020 | B.1.83 | May | 18 |
| 02/05/2020 | B.1.1 | May | 18 |
| 02/05/2020 | B.1.1.1 | May | 18 |
| 02/05/2020 | B.1.1 | May | 18 |

| | | | |
|---|---|---|---|
| 02/05/2020 | B.1.1.1 | May | 18 |
| 03/05/2020 | B.1.83 | May | 18 |
| 03/05/2020 | B.1.1.5 | May | 18 |
| 03/05/2020 | B.1.1.1 | May | 18 |
| 05/05/2020 | B.1.5 | May | 18 |
| 06/05/2020 | B.1.1.5 | May | 19 |
| 06/05/2020 | B.1.1 | May | 19 |
| 07/05/2020 | B.1.6 | May | 19 |
| 07/05/2020 | B.1.1.75 | May | 19 |
| 07/05/2020 | B.1 | May | 19 |
| 08/05/2020 | B.1.1 | May | 19 |
| 08/05/2020 | B.1.83 | May | 19 |
| 08/05/2020 | B.1.94 | May | 19 |
| 08/05/2020 | B.1.1.1 | May | 19 |
| 08/05/2020 | B.1.1.1 | May | 19 |
| 08/05/2020 | B.1.1 | May | 19 |
| 08/05/2020 | B.1.1 | May | 19 |
| 08/05/2020 | B.1.1 | May | 19 |
| 09/05/2020 | B.1.1.1 | May | 19 |
| 09/05/2020 | B.1.83 | May | 19 |
| 09/05/2020 | B.1.1 | May | 19 |
| 09/05/2020 | B.1.1 | May | 19 |
| 09/05/2020 | B.1.5 | May | 19 |
| 09/05/2020 | B.1.83 | May | 19 |
| 10/05/2020 | B.1.1.1 | May | 19 |
| 12/05/2020 | B.1 | May | 19 |
| 12/05/2020 | B.1 | May | 19 |
| 12/05/2020 | B.1.1.1 | May | 19 |
| 13/05/2020 | B.1.1 | May | 20 |
| 13/05/2020 | B.1.1 | May | 20 |
| 13/05/2020 | B.1.1 | May | 20 |
| 14/05/2020 | B.1.1 | May | 20 |
| 14/05/2020 | B.1.1 | May | 20 |
| 14/05/2020 | B.1.1.5 | May | 20 |
| 14/05/2020 | B.1.1 | May | 20 |
| 16/05/2020 | B.1.1.1 | May | 20 |
| 17/04/2020 | B.1.1.1 | Apr | 16 |
| 17/05/2020 | B.1.1.5 | May | 20 |
| 17/05/2020 | B.1.1.1 | May | 20 |
| 17/05/2020 | B.1.1.1 | May | 20 |
| 21/05/2020 | B.1.1 | May | 21 |
| 21/05/2020 | B.1 | May | 21 |
| 21/05/2020 | B.1.1.5 | May | 21 |
| 21/05/2020 | B.1.1.1 | May | 21 |
| 26/05/2020 | B.1.1 | May | 21 |
| 28/05/2020 | B.1.1.1 | May | 22 |

| | | | |
|---|---|---|---|
| 28/05/2020 | B.1.1.1 | May | 22 |
| 02/06/2020 | B.1 | Jun | 22 |
| 03/06/2020 | B.1.1 | Jun | 23 |
| 10/06/2020 | B.1 | Jun | 24 |
| 11/06/2020 | B.1.1.221 | Jun | 24 |
| 11/06/2020 | B.1.256 | Jun | 24 |
| 12/06/2020 | B.1.1.221 | Jun | 24 |
| 15/06/2020 | B.1.1.221 | Jun | 24 |
| 15/06/2020 | B.1.1.221 | Jun | 24 |
| 15/06/2020 | B.1.1.221 | Jun | 24 |
| 15/06/2020 | B.1.1 | Jun | 24 |
| 13/03/2020 | B.1 | Mar | 11 |
| 16/05/2020 | B.1.5 | May | 20 |
| 02/03/2020 | B.1.5 | Mar | 9 |
| 03/03/2020 | B.1.1 | Mar | 9 |
| 03/03/2020 | B.1 | Mar | 9 |
| 04/03/2020 | B.1.1 | Mar | 10 |
| 05/03/2020 | B.1.5 | Mar | 10 |
| 06/03/2020 | B.1 | Mar | 10 |
| 07/03/2020 | B.1.1 | Mar | 10 |
| 07/03/2020 | B.1.1 | Mar | 10 |
| 12/03/2020 | B.29 | Mar | 11 |
| 13/03/2020 | B | Mar | 11 |
| 09/04/2020 | B.1.8 | Apr | 15 |
| 09/04/2020 | B | Apr | 15 |
| 11/04/2020 | B.1.5 | Apr | 15 |
| 14/04/2020 | B.1.1.5 | Apr | 15 |
| 14/04/2020 | B.1.157 | Apr | 15 |
| 14/04/2020 | B.1.5 | Apr | 15 |
| 08/05/2020 | B.1.1.1 | May | 19 |
| 08/05/2020 | B.1.1 | May | 19 |
| 12/05/2020 | B.1.1 | May | 19 |
| 21/05/2020 | B.1.5 | May | 21 |
| 04/06/2020 | B.1.1 | Jun | 23 |
| 07/06/2020 | B.1.1 | Jun | 23 |

Suppl. File 1. GISAID genome accession numbers from SARS-CoV2, Belgium (February 2020 – June 2020).

EPI_ISL_407976

EPI_ISL_415159

EPI_ISL_415153

EPI_ISL_415154

EPI_ISL_415155

EPI_ISL_415156

EPI_ISL_415158

EPI_ISL_415157

EPI_ISL_416467

EPI_ISL_416468

EPI_ISL_416470

EPI_ISL_416471

EPI_ISL_416472

EPI_ISL_416476

EPI_ISL_417427

EPI_ISL_417428

EPI_ISL_418986

EPI_ISL_738198

EPI_ISL_416469

EPI_ISL_416475

EPI_ISL_417429

EPI_ISL_417430

EPI_ISL_418987

EPI_ISL_734503

EPI_ISL_734489

EPI_ISL_738199

EPI_ISL_738200

EPI_ISL_417422

EPI_ISL_417424

EPI_ISL_417426

EPI_ISL_418270

EPI_ISL_418988

EPI_ISL_418989

EPI_ISL_420442

EPI_ISL_420443

EPI_ISL_420314

EPI_ISL_420447

EPI_ISL_420448

EPI_ISL_420319

EPI_ISL_420454

EPI_ISL_420324

EPI_ISL_738201

EPI_ISL_418863

EPI_ISL_418792

EPI_ISL_418796

EPI_ISL_418798

EPI_ISL_418806

EPI_ISL_420444

EPI_ISL_420317

EPI_ISL_420453

EPI_ISL_420322

EPI_ISL_420325

EPI_ISL_420329

EPI_ISL_420330

EPI_ISL_420331

EPI_ISL_420334

EPI_ISL_420337

EPI_ISL_420339

EPI_ISL_420344

EPI_ISL_420348

EPI_ISL_420349

EPI_ISL_420353

EPI_ISL_420354

EPI_ISL_420367

EPI_ISL_738202

EPI_ISL_417425

EPI_ISL_418981

EPI_ISL_418794

EPI_ISL_418795

EPI_ISL_418805

EPI_ISL_420315

EPI_ISL_420446

EPI_ISL_420449

EPI_ISL_420441

EPI_ISL_420321

EPI_ISL_420326

EPI_ISL_420328

EPI_ISL_420332

EPI_ISL_420333

EPI_ISL_420335

EPI_ISL_420336

EPI_ISL_420340

EPI_ISL_420341

EPI_ISL_420342

EPI_ISL_420343

EPI_ISL_420345

EPI_ISL_420347

EPI_ISL_420350

EPI_ISL_420351

EPI_ISL_420355

EPI_ISL_420356

EPI_ISL_738203

EPI_ISL_420316

EPI_ISL_420450

EPI_ISL_420318

EPI_ISL_420451

EPI_ISL_420323

EPI_ISL_420327

EPI_ISL_420352

EPI_ISL_738204

EPI_ISL_738205

EPI_ISL_418982

EPI_ISL_418983

EPI_ISL_418984

EPI_ISL_418985

EPI_ISL_420445

EPI_ISL_420320

EPI_ISL_420452

EPI_ISL_420338

EPI_ISL_420346

EPI_ISL_522349

EPI_ISL_418793

EPI_ISL_418800

EPI_ISL_462210

EPI_ISL_734485

EPI_ISL_462255

EPI_ISL_734504

EPI_ISL_738206

EPI_ISL_738196

EPI_ISL_738207

EPI_ISL_734505

EPI_ISL_458230

EPI_ISL_420361

EPI_ISL_462178

EPI_ISL_462179

EPI_ISL_462181

EPI_ISL_734506

EPI_ISL_420358

EPI_ISL_420359

EPI_ISL_420360

EPI_ISL_420362

EPI_ISL_420364

EPI_ISL_420366

EPI_ISL_420368

EPI_ISL_420369

EPI_ISL_420370

EPI_ISL_458229

EPI_ISL_458235

EPI_ISL_734507

EPI_ISL_734508

EPI_ISL_734509

EPI_ISL_420357

EPI_ISL_420363

EPI_ISL_458176

EPI_ISL_458228

EPI_ISL_458232

EPI_ISL_462162

EPI_ISL_734510

EPI_ISL_734511

EPI_ISL_734512

EPI_ISL_734513

EPI_ISL_462209

EPI_ISL_462265

EPI_ISL_734514

EPI_ISL_734515

EPI_ISL_734516

EPI_ISL_420371

EPI_ISL_420372

EPI_ISL_420373

EPI_ISL_420386

EPI_ISL_420396

EPI_ISL_420397

EPI_ISL_420398

EPI_ISL_420411

EPI_ISL_420416

EPI_ISL_420432

EPI_ISL_462263

EPI_ISL_462264

EPI_ISL_734517

EPI_ISL_734518

EPI_ISL_734519

EPI_ISL_420374

EPI_ISL_420375

EPI_ISL_420376

EPI_ISL_420377

EPI_ISL_420378

EPI_ISL_420381

EPI_ISL_420382

EPI_ISL_420383

EPI_ISL_420387

EPI_ISL_420388

EPI_ISL_420389

EPI_ISL_420390

EPI_ISL_420391

EPI_ISL_420392

EPI_ISL_420393

EPI_ISL_420395

EPI_ISL_420399

EPI_ISL_420401

EPI_ISL_420406

EPI_ISL_420407

EPI_ISL_420408

EPI_ISL_420412

EPI_ISL_420417

EPI_ISL_420418

EPI_ISL_420420

EPI_ISL_420421

EPI_ISL_420424

EPI_ISL_420425

EPI_ISL_420426

EPI_ISL_420427

EPI_ISL_420429

EPI_ISL_420430

EPI_ISL_420431

EPI_ISL_420433

EPI_ISL_420434

EPI_ISL_420436

EPI_ISL_462187

EPI_ISL_734520

EPI_ISL_420379

EPI_ISL_420380

EPI_ISL_420384

EPI_ISL_420385

EPI_ISL_420394

EPI_ISL_420400

EPI_ISL_420402

EPI_ISL_420403

EPI_ISL_420404

EPI_ISL_420405

EPI_ISL_420409

EPI_ISL_420410

EPI_ISL_420413

EPI_ISL_420414

EPI_ISL_420415

EPI_ISL_420419

EPI_ISL_420422

EPI_ISL_420423

EPI_ISL_420428

EPI_ISL_420435

EPI_ISL_420437

EPI_ISL_420438

EPI_ISL_420439

EPI_ISL_420440

EPI_ISL_462188

EPI_ISL_462191

EPI_ISL_462193

EPI_ISL_462194

EPI_ISL_462234

EPI_ISL_462256

EPI_ISL_734521

EPI_ISL_734522

EPI_ISL_458231

EPI_ISL_462169

EPI_ISL_462170

EPI_ISL_462176

EPI_ISL_462177

EPI_ISL_462190

EPI_ISL_462204

EPI_ISL_462245

EPI_ISL_734523

EPI_ISL_458180

EPI_ISL_458181

EPI_ISL_458182

EPI_ISL_458183

EPI_ISL_458184

EPI_ISL_458185

EPI_ISL_458186

EPI_ISL_462171

EPI_ISL_462172

EPI_ISL_462173

EPI_ISL_462174

EPI_ISL_462175

EPI_ISL_462198

EPI_ISL_462199

EPI_ISL_462200

EPI_ISL_462202

EPI_ISL_462203

EPI_ISL_462207

EPI_ISL_462223

EPI_ISL_462244

EPI_ISL_458156

EPI_ISL_458157

EPI_ISL_458158

EPI_ISL_458159

EPI_ISL_458160

EPI_ISL_458161

EPI_ISL_458162

EPI_ISL_458163

EPI_ISL_458164

EPI_ISL_458165

EPI_ISL_458166

EPI_ISL_458167

EPI_ISL_458168

EPI_ISL_458169

EPI_ISL_458170

EPI_ISL_458171

EPI_ISL_458172

EPI_ISL_458173

EPI_ISL_458174

EPI_ISL_458175

EPI_ISL_458177

EPI_ISL_458178

EPI_ISL_458179

EPI_ISL_462163

EPI_ISL_462164

EPI_ISL_462165

EPI_ISL_462166

EPI_ISL_462167

EPI_ISL_462168

EPI_ISL_462192

EPI_ISL_462195

EPI_ISL_462196

EPI_ISL_462197

EPI_ISL_462208

EPI_ISL_462224

EPI_ISL_462242

EPI_ISL_734524

EPI_ISL_458187

EPI_ISL_458188

EPI_ISL_458189

EPI_ISL_458190

EPI_ISL_458191

EPI_ISL_458192

EPI_ISL_458193

EPI_ISL_458194

EPI_ISL_458195

EPI_ISL_458196

EPI_ISL_458197

EPI_ISL_458198

EPI_ISL_458199

EPI_ISL_458200

EPI_ISL_458201

EPI_ISL_458202

EPI_ISL_458203

EPI_ISL_458204

EPI_ISL_458205

EPI_ISL_458206

EPI_ISL_458207

EPI_ISL_458208

EPI_ISL_458209

EPI_ISL_458210

EPI_ISL_458211

EPI_ISL_458212

EPI_ISL_458213

EPI_ISL_458214

EPI_ISL_458215

EPI_ISL_458216

EPI_ISL_462206

EPI_ISL_462211

EPI_ISL_462240

EPI_ISL_462243

EPI_ISL_462246

EPI_ISL_734486

EPI_ISL_458217

EPI_ISL_458218

EPI_ISL_458219

EPI_ISL_458220

EPI_ISL_458221

EPI_ISL_458222

EPI_ISL_458223

EPI_ISL_458224

EPI_ISL_458225

EPI_ISL_458226

EPI_ISL_458227

EPI_ISL_462158

EPI_ISL_462159

EPI_ISL_462160

EPI_ISL_462161

EPI_ISL_462182

EPI_ISL_462183

EPI_ISL_462184

EPI_ISL_462185

EPI_ISL_462186

EPI_ISL_462189

EPI_ISL_462201

EPI_ISL_462205

EPI_ISL_462212

EPI_ISL_462217

EPI_ISL_462219

EPI_ISL_462221

EPI_ISL_462238

EPI_ISL_462239

EPI_ISL_462241

EPI_ISL_734525

EPI_ISL_458233

EPI_ISL_458234

EPI_ISL_462213

EPI_ISL_462214

EPI_ISL_462215

EPI_ISL_462216

EPI_ISL_462231

EPI_ISL_462232

EPI_ISL_462233

EPI_ISL_734526

EPI_ISL_734527

EPI_ISL_462218

EPI_ISL_462220

EPI_ISL_462222

EPI_ISL_462225

EPI_ISL_462226

EPI_ISL_462227

EPI_ISL_462229

EPI_ISL_462230

EPI_ISL_734528

EPI_ISL_734529

EPI_ISL_734530

EPI_ISL_462228

EPI_ISL_462237

EPI_ISL_734531

EPI_ISL_464084

EPI_ISL_462235

EPI_ISL_462236

EPI_ISL_734532

EPI_ISL_734533

EPI_ISL_734534

EPI_ISL_464070

EPI_ISL_464071

EPI_ISL_734535

EPI_ISL_464083

EPI_ISL_464067

EPI_ISL_464069

EPI_ISL_464072

EPI_ISL_464073

EPI_ISL_464079

EPI_ISL_462252

EPI_ISL_462254

EPI_ISL_734536

EPI_ISL_734537

EPI_ISL_734538

EPI_ISL_734539

EPI_ISL_464065

EPI_ISL_464066

EPI_ISL_464068

EPI_ISL_464074

EPI_ISL_464076

EPI_ISL_464077

EPI_ISL_464081

EPI_ISL_464082

EPI_ISL_462247

EPI_ISL_462248

EPI_ISL_462249

EPI_ISL_462250

EPI_ISL_462251

EPI_ISL_462257

EPI_ISL_462258

EPI_ISL_462259

EPI_ISL_734540

EPI_ISL_734541

EPI_ISL_734542

EPI_ISL_734543

EPI_ISL_464075

EPI_ISL_464078

EPI_ISL_464080

EPI_ISL_464086

EPI_ISL_464087

EPI_ISL_462253

EPI_ISL_462266

EPI_ISL_462267

EPI_ISL_462268

EPI_ISL_734544

EPI_ISL_734545

EPI_ISL_734546

EPI_ISL_734547

EPI_ISL_734548

EPI_ISL_734549

EPI_ISL_464085

EPI_ISL_464088

EPI_ISL_464089

EPI_ISL_464090

EPI_ISL_462260

EPI_ISL_462261

EPI_ISL_462269

EPI_ISL_476941

EPI_ISL_476949

EPI_ISL_476942

EPI_ISL_476943

EPI_ISL_476944

EPI_ISL_476945

EPI_ISL_476946

EPI_ISL_476947

EPI_ISL_476948

EPI_ISL_734550

EPI_ISL_734551

EPI_ISL_734552

EPI_ISL_734553

EPI_ISL_734554

EPI_ISL_734555

EPI_ISL_734556

EPI_ISL_734557

EPI_ISL_734558

EPI_ISL_734559

EPI_ISL_738208

EPI_ISL_738209

EPI_ISL_734560

EPI_ISL_734561

EPI_ISL_734562

EPI_ISL_734563

EPI_ISL_734564

EPI_ISL_462262

EPI_ISL_476950

EPI_ISL_476951

EPI_ISL_476952

EPI_ISL_476953

EPI_ISL_476954

EPI_ISL_476955

EPI_ISL_476956

EPI_ISL_476957

EPI_ISL_476958

EPI_ISL_476959

EPI_ISL_476960

EPI_ISL_476961

EPI_ISL_476962

EPI_ISL_476963

EPI_ISL_476964

EPI_ISL_476965

EPI_ISL_476966

EPI_ISL_476967

EPI_ISL_476968

EPI_ISL_476969

EPI_ISL_476970

EPI_ISL_476971

EPI_ISL_476972

EPI_ISL_476973

EPI_ISL_476974

EPI_ISL_476975

EPI_ISL_476976

EPI_ISL_476977

EPI_ISL_476978

EPI_ISL_476979

EPI_ISL_476980

EPI_ISL_476981

EPI_ISL_476982

EPI_ISL_738210

EPI_ISL_734565

EPI_ISL_734566

EPI_ISL_734567

EPI_ISL_734568

EPI_ISL_734569

EPI_ISL_734570

EPI_ISL_734571

EPI_ISL_476983

EPI_ISL_476984

EPI_ISL_476985

EPI_ISL_476986

EPI_ISL_476987

EPI_ISL_476988

EPI_ISL_476989

EPI_ISL_734572

EPI_ISL_734573

EPI_ISL_734574

EPI_ISL_734575

EPI_ISL_734576

EPI_ISL_734577

EPI_ISL_734578

EPI_ISL_734579

EPI_ISL_734580

EPI_ISL_734581

EPI_ISL_734582

EPI_ISL_738211

EPI_ISL_738212

EPI_ISL_738213

EPI_ISL_734583

EPI_ISL_734584

EPI_ISL_734585

EPI_ISL_734586

EPI_ISL_734587

EPI_ISL_734588

EPI_ISL_734589

EPI_ISL_734590

EPI_ISL_734591

EPI_ISL_734592

EPI_ISL_734593

EPI_ISL_734594

EPI_ISL_734595

EPI_ISL_734596

EPI_ISL_734597

EPI_ISL_734598

EPI_ISL_734599

EPI_ISL_734600

EPI_ISL_734601

EPI_ISL_734602

EPI_ISL_734603

EPI_ISL_734604

EPI_ISL_734605

EPI_ISL_734606

EPI_ISL_734607

EPI_ISL_734608

EPI_ISL_734609

EPI_ISL_734610

EPI_ISL_734611

EPI_ISL_734612

EPI_ISL_734613

EPI_ISL_734736

EPI_ISL_734614

EPI_ISL_734615

EPI_ISL_734616

EPI_ISL_734617

EPI_ISL_734618

EPI_ISL_734619

EPI_ISL_476990

EPI_ISL_734620

EPI_ISL_734621

EPI_ISL_734622

EPI_ISL_734623

EPI_ISL_734624

EPI_ISL_734625

EPI_ISL_734626

EPI_ISL_734627

EPI_ISL_734628

EPI_ISL_734629

EPI_ISL_734630

EPI_ISL_734631

EPI_ISL_734632

EPI_ISL_476991

EPI_ISL_734633

EPI_ISL_734634

EPI_ISL_734635

EPI_ISL_734636

EPI_ISL_734637

EPI_ISL_734638

EPI_ISL_734639

EPI_ISL_462151

EPI_ISL_462152

EPI_ISL_462153

EPI_ISL_462154

EPI_ISL_462155

EPI_ISL_462156

EPI_ISL_462157

EPI_ISL_734640

EPI_ISL_734641

EPI_ISL_734642

EPI_ISL_734643

EPI_ISL_734644

EPI_ISL_734645

EPI_ISL_734646

EPI_ISL_734647

EPI_ISL_734648

EPI_ISL_734649

EPI_ISL_734653

EPI_ISL_734654

EPI_ISL_734655

EPI_ISL_734656

EPI_ISL_734650

EPI_ISL_734651

EPI_ISL_734652

EPI_ISL_476992

EPI_ISL_734658

EPI_ISL_734659

EPI_ISL_734660

EPI_ISL_734661

EPI_ISL_734662

EPI_ISL_734663

EPI_ISL_734664

EPI_ISL_734665

EPI_ISL_734666

EPI_ISL_734667

EPI_ISL_734668

EPI_ISL_734669

EPI_ISL_734657

EPI_ISL_476993

EPI_ISL_476994

EPI_ISL_734670

EPI_ISL_476995

EPI_ISL_734671

EPI_ISL_734672

EPI_ISL_734673

EPI_ISL_734674

EPI_ISL_734675

EPI_ISL_734676

EPI_ISL_734677

EPI_ISL_734678

EPI_ISL_734679

EPI_ISL_734680

EPI_ISL_734681

EPI_ISL_734500

EPI_ISL_734682

EPI_ISL_734683

EPI_ISL_734685

EPI_ISL_734684

EPI_ISL_734686

EPI_ISL_734687

EPI_ISL_734688

EPI_ISL_476996

EPI_ISL_734689

EPI_ISL_734690

EPI_ISL_734691

EPI_ISL_734692

EPI_ISL_734693

EPI_ISL_734694

EPI_ISL_476997

EPI_ISL_476998

EPI_ISL_734695

EPI_ISL_734696

EPI_ISL_734697

EPI_ISL_734698

EPI_ISL_734699

EPI_ISL_734700

EPI_ISL_476999

EPI_ISL_734701

EPI_ISL_734702

EPI_ISL_734703

EPI_ISL_477000

EPI_ISL_477001

EPI_ISL_477002

EPI_ISL_734704

EPI_ISL_462270

EPI_ISL_477004

EPI_ISL_734705

EPI_ISL_734706

EPI_ISL_462271

EPI_ISL_462272

EPI_ISL_462273

EPI_ISL_462274

EPI_ISL_462275

EPI_ISL_477005

EPI_ISL_734707

EPI_ISL_734708

EPI_ISL_734709

EPI_ISL_734710

EPI_ISL_734711

EPI_ISL_734712

EPI_ISL_734713

EPI_ISL_734714

EPI_ISL_734715

EPI_ISL_734716

EPI_ISL_734717

EPI_ISL_738214

EPI_ISL_738215

EPI_ISL_734718

EPI_ISL_734719

EPI_ISL_734720

EPI_ISL_734721

EPI_ISL_734722

EPI_ISL_734723

EPI_ISL_734724

EPI_ISL_477006

EPI_ISL_734492

EPI_ISL_734725

EPI_ISL_734726

EPI_ISL_734727

EPI_ISL_738216

EPI_ISL_477007

EPI_ISL_734728

EPI_ISL_734729

EPI_ISL_734730

EPI_ISL_734731

EPI_ISL_734732

EPI_ISL_734733

EPI_ISL_734734

EPI_ISL_734735

EPI_ISL_738197

EPI_ISL_734737

EPI_ISL_734738

EPI_ISL_734739

EPI_ISL_734740

EPI_ISL_734741

EPI_ISL_734742

EPI_ISL_734743

EPI_ISL_738217

EPI_ISL_734744

EPI_ISL_734745

EPI_ISL_734746

EPI_ISL_734747

EPI_ISL_734748

EPI_ISL_738218

EPI_ISL_738219

EPI_ISL_522350

EPI_ISL_734749

EPI_ISL_734750

EPI_ISL_734751

EPI_ISL_734493

EPI_ISL_734752

EPI_ISL_734753

EPI_ISL_734754

EPI_ISL_734755

EPI_ISL_734756
